# Targeted Phasing of 2-200 Kilobase DNA Fragments with a Short-Read Sequencer and a Single-Tube Linked-Read Library Method

**DOI:** 10.1101/2023.03.05.531179

**Authors:** Veronika Mikhaylova, Madison Rzepka, Tetsuya Kawamura, Yu Xia, Peter L. Chang, Shiguo Zhou, Long Pham, Naisarg Modi, Likun Yao, Adrian Perez-Agustin, Sara Pagans, T. Christian Boles, Ming Lei, Yong Wang, Ivan Garcia-Bassets, Zhoutao Chen

## Abstract

In the human genome, heterozygous sites are genomic positions with different alleles inherited from each parent. On average, there is a heterozygous site every 1-2 kilobases (kb). Resolving whether two alleles in neighboring heterozygous positions are physically linked—that is, phased—is possible with a short-read sequencer if the sequencing library captures long-range information. TELL-Seq is a library preparation method based on millions of barcoded micro-sized beads that enables instrument-free phasing of a whole human genome in a single PCR tube. TELL-Seq incorporates a unique molecular identifier (barcode) to the short reads generated from the same high-molecular-weight (HMW) DNA fragment (known as ‘linked-reads’). However, genome-scale TELL-Seq is not cost-effective for applications focusing on a single locus or a few loci. Here, we present an optimized TELL-Seq protocol that enables the cost-effective phasing of enriched loci (targets) of varying sizes, purity levels, and heterozygosity. Targeted TELL-Seq maximizes linked-read efficiency and library yield while minimizing input requirements, fragment collisions on microbeads, and sequencing burden. To validate the targeted protocol, we phased seven 180-200 kb loci enriched by CRISPR/Cas9-mediated excision coupled with pulse-field electrophoresis, four 20 kb loci enriched by CRISPR/Cas9-mediated protection from exonuclease digestion, and six 2-13 kb loci amplified by PCR. The selected targets have clinical and research relevance (*BRCA1, BRCA2, MLH1, MSH2, MSH6, APC, PMS2, SCN5A*-*SCN10A*, and *PKI3CA*). These analyses reveal that targeted TELL-Seq provides a reliable way of phasing allelic variants within targets (2-200 kb in length) with the low cost and high accuracy of short-read sequencing.

## INTRODUCTION

Polymorphic sites are genomic positions with two or more nucleotide or length variants across and within human populations ^1^. In an individual human genome, a polymorphic site can carry a different variant (‘allele’) inherited from each parent (known as ‘heterozygous site’). Resolving (‘phasing’) these sequence differences by parental copy is critical to untangling human evolution and understanding the genetic basis of complex traits and diseases ^2,3^. On average, humans carry a heterozygosity every 1-2 kilobases (kb), depending on genetic ancestry and admixture ^4–8^; thus, sequencing platforms such as Pacific Biosciences^®^ (PacBio) and Oxford Nanopore Technology^®^ (ONT) that process 20-100 kb long reads have drawn much attention ^9^. However, long-read sequencers remain less cost-effective and can have lower genotyping fidelity and throughput than short-read sequencers, especially for reads longer than 20 kb ^10^. Moreover, newer short-read technologies are evolving and further pushing down sequencing costs (e.g., Ultima Genomics^®^, Singular Genomics^®^, and Element Biosciences^®^). Therefore, incorporating long-range information into short reads remains an attractive option and will likely remain as such if short-read sequencing costs soon drop to a hundred-dollar per genome.

Long-range genomic information can be incorporated into short reads through DNA co-barcoding reactions (i.e., two or more barcoding events on the same DNA molecule). DNA co-barcoding relies on tagging subfragments derived from individual HMW DNA molecules with unique molecular identifiers (barcodes). On a high throughput scale, millions of paralleled DNA co-barcoding reactions enable the phasing of a whole human genome, which can be achieved with two different molecular strategies: synthetic long reads and linked reads. Synthetic long reads enable co-barcoding DNA fragment length within ∼6-10 kb distances and require over-sequencing for best results ^11–13^, while linked reads enable co-barcoding DNA fragments with much longer lengths (50-200 kb), reducing the need for over-sequencing ^14^. There are three subcategories of linked-read methods: droplet-based ^15–20^, microwell-based ^14,21,22^, and microbead-based ^23,24^. Microbead-based linked-read methods are also referred to as ‘single-tube’ as they enable phasing in a single-tube reaction. TELL-Seq™ (Transposase Enzyme Linked Long-read Sequencing) is a single-tube linked-read method with optimized chemistry that facilitates simultaneous co-barcoding reactions on the surface of millions of microbeads ^24^. TELL-Seq libraries can be generated in only three hours and sequenced on a short-read instrument leveraging the low cost, broad accessibility, and accuracy of these platforms ^24^.

Genome-scale phasing is not cost-effective for clinical and research applications focusing on a single locus or a few loci (targeted approaches). To save on sequencing burden and computational running time, a step of DNA enrichment can be added before sequencing library preparation. There are three classes of DNA pre-enrichment methods: probe-based by hybridization ^25,26^; CRISPR/Cas9-based by RNA-guided Cas9 nuclease-mediated excision ^27^; and long-range PCR ^18^. In principle, the three methods should be compatible with TELL-Seq, but the TELL-Seq protocol may require adaptation to remain cost-effective and provide high-quality phased data. For example, genome-scale TELL-Seq is permissive to multiple DNA molecules (collisions) on the surface of the same microbead (an average of 6-8) since it is unlikely that two or more of these molecules are copies of the same locus ^24^. However, a high collision rate would be detrimental when processing a single target due to the ultra-low complexity of this sample. Additionally, genome-scale phasing requires a high number of barcodes to phase a whole genome ^24^, whereas targeted phasing requires a much lower number of barcodes owing to the ultra-low complexity of a single locus. Here, we developed and validated TELL-Seq conditions that can generate high-quality and cost-effective phased data with a variety of target sizes, off-target background levels, and heterozygosity.

## RESULTS

### Adapting TELL-Seq to a targeted scale

TELL-Seq was initially developed to capture long-range information from whole genomes (human, animal, plant, invertebrate, or microbe) or metagenomes—also known as WGS TELL-Seq ^24^. However, capturing long-range information from a single locus or a few loci, hereafter referred to as targeted TELL-Seq, presents some challenges. We capitalized upon our experience in adapting WGS TELL-Seq to genomes of different sizes to devise a series of protocol adjustments that should enable high-quality phased data with target inputs. The key elements of a targeted TELL-Seq protocol would be using ≤100 pg input DNA and approximately 2 million (6 μL) TELL microbeads in a 25 μL reaction during the barcoding step and using no more than 250,000 target-bound TELL microbeads (≤0.75 μL) in a 25 μL PCR reaction volume and amplifying with up to 21 cycles during the library amplification step. Compare WGS TELL-Seq protocols for human and microbe in **Suppl. Fig. 1A** and other genomes in **Suppl. Fig. 1B** (WGS column), with targeted TELL-Seq options in **Suppl. Fig. 1B**, (column with text in red). The indicated volumes, amounts, and number of PCR cycles in the proposed targeted TELL-Seq protocol have been designed to maximize the linked-read efficiency and library yield while minimizing DNA collisions on microbeads and sequencing burden.

### Phasing partially enriched, ultra-long targets

To validate the targeted TELL-Seq protocol, we first focused on a set of ultra-long targets (∼200 kb) excised from genomic DNA using the HLS-CATCH method ^27^. The HLS-CATCH workflow preserves ultra-long HMW DNA (up to 1 Mb) by integrating cell lysis, genomic DNA extraction, and target isolation in the same platform, avoiding DNA shearing from liquid handling and centrifugation steps. In CATCH process, targets are excised from genomic DNA using guide (g)RNA-directed Cas9 ribonucleoprotein (RNP) complexes and isolated using preparative pulse-field electrophoresis (**Suppl. Fig. 2A** and **Fig. 1A**) ^27–30^. After enrichment, targets typically represent a 0.5-5% of the total DNA—hereafter referred to as ‘partially enriched targets’—which can still represent more than a 150-fold enrichment over non-target DNA of similar size (**Suppl. Fig. 1B**, Targeted, 0.5-5% option). This type of impure sample will still be relatively permissive to DNA collisions owing to the complexity of the non-target background. As targets, we selected a group of genomic loci with recurrent germline mutations associated with hereditary predisposition to cancer: *BRCA1* (target length: 198.4 kb), *BRCA2* (187.5 kb), *APC* (199.6 kb), *MSH2* (201.1 kb), *MSH6* (197.5 kb), *MLH1* (201.7 kb), and *PMS2* (197.5 kb).

**Figure 1.**
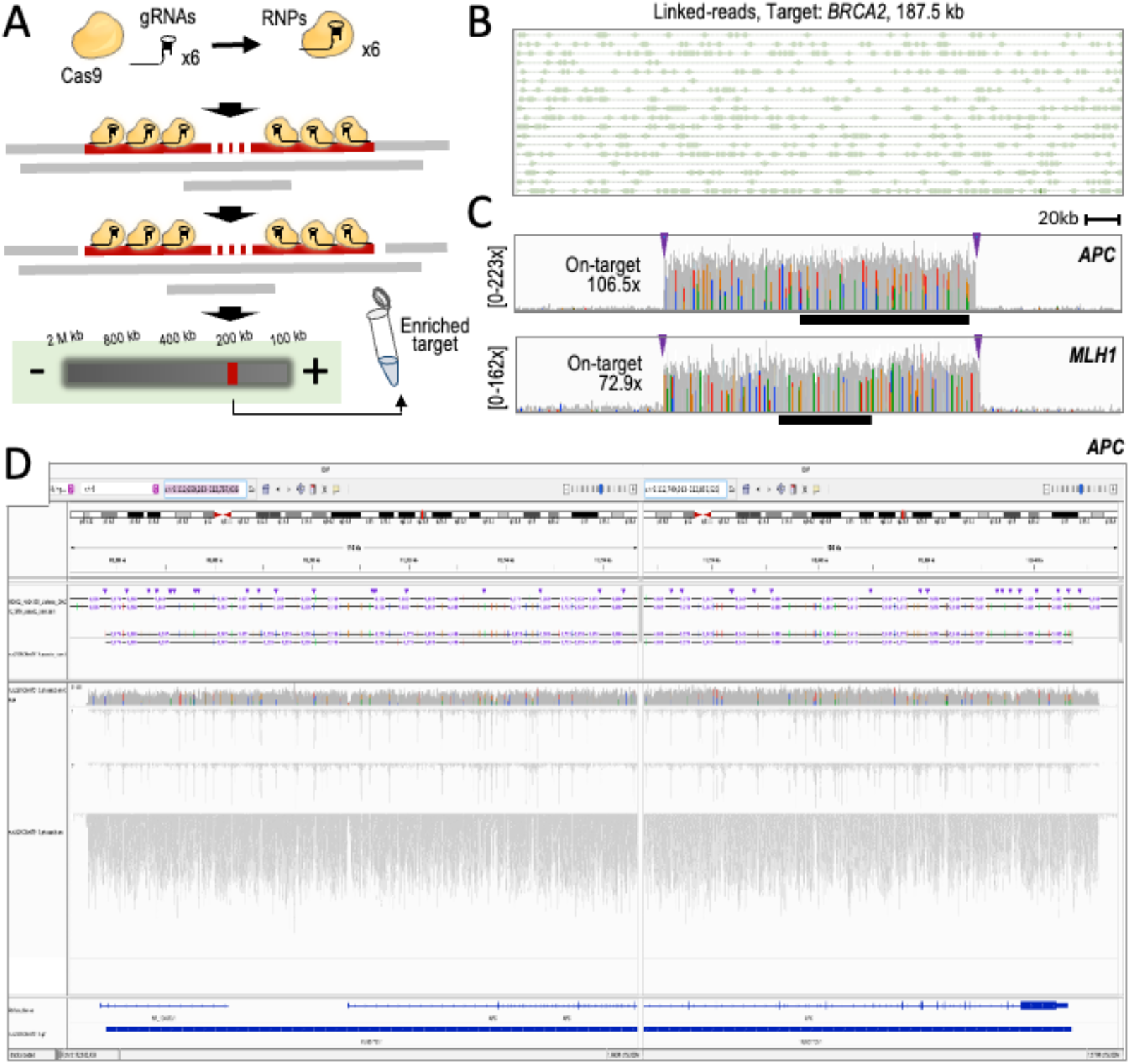
Phasing of -180-200 kb targets enriched with the HLS-CATCH system. **A**. Experimental workflow (see also Suppl. Fig. 1A). **B**. A representative example of linked reads obtained with Targeted TELL-Seq. Shown a 186 kb region in the *BRCA2* target. Lines (linked reads) join uniquely barcoded short reads (boxes). **C**. Read profiles (coverage indicated) from TELL-Seq libraries showing robust on-target recovery compared to background. Targets: 200 kb *APC* and *MLH1* targets. Arrowheads represent locations of 3x gRNA binding sites. Black bars represent the *APC* and *MLH1* genes. **D**. Joint screenshots from the IGV Portal showing TELL-Seq results for the *APC* target. Tracks (from top to bottom): GIAB phased haplotypes in HG002 (benchmark); phased haplotypes (numbers indicate distances between some phased sites; coverage; inferred haplotype 1; inferred haplotype 2; unphased reads; gene annotations; and inferred phase block.

We designed three gRNAs complementary to each end of every target in order to maximize target excision and minimize the risk of haplotype dropout due to target-specific allelic differences at the Grna binding site (**Fig. 1A**). We then assembled Cas9 RNP complexes in three distinct pools: a RNP pool with six *BRCA1* gRNAs, another RNP pool with six *BRCA2* gRNAs, and a third RNP pool with thirty gRNAs for *APC, MSH2, MSH6, MLH1*, and *PMS2*. We used GM24149 and GM24385 B lymphocytes from Genome in a Bottle (GIAB) as sources of genomic DNA, which are two well-annotated reference materials: HG003 (NA24149) and HG002 (NA24385), respectively. Interestingly, our use of HG002 added two singularities to the planned phasing tests. The first singularity is the relatively low density of heterozygous sites in the HG002 *MLH1* locus with a median distance between adjacent sites of >13 kb, which is more than nineteen times the median distance between heterozygous sites observed in the other six targets (**Suppl. Fig. 2B**). The second singularity is the presence of a large polymorphic inversion relocating the three 3’ gRNA binding sites more than 700 bp downstream from the expected positions in the HG002 *PMS2* locus (**Suppl. Fig. 2C**, see scheme and red arrowheads at the bottom of the panel) ^31,32^. We used GIAB-generated WGS linked-read data from HG002 to confirm the presence of this inversion (**Suppl. Fig. 2C**, left heatmaps show 5.4 Mb regions while the right heatmap shows a 1.35 Mb region). Besides testing the TELL-Seq protocol with typical targets, we aimed to better understand the process of targeted phasing with challenging targets.

We used HG003 as a source of genomic DNA to isolate the *BRCA1* and *BRCA2* targets and HG002 as a source of genomic DNA to isolate the other five targets as a pool. We recovered the three excised DNA fractions of around ∼200 kb from the HLS-CATCH system. We then mixed 100 pg of the *BRCA1* and *BRCA2* targets to generate a single TELL-Seq library and used 270 pg of the multi-loci mixture to generate a second TELL-Seq library. After sequencing and duplicate removal, we obtained 10.4 million and 23.1 million reads from the two libraries, respectively. The mapping profiles were consistent with those found in conventional WGS TELL-Seq (**Fig. 1B**). On-target read recovery was 3.7% for *BRCA1+BRCA2* and 5.4% for the other five targets, with an average mean coverage of 85.4x and 108.5x, respectively (**Fig. 1C** shows read profiles for two representative cases). The lowest recovery was observed for the two targets with genomic singularities, 0.9%/72.9x (*MLH1*) and 0.4%/36.1x (*PMS2*), compared to 1.9%/106.4x (*BRCA1*), 1.8%/108.5x (*BRCA2*), 1.3%/106.5x (*APC*), 1.4%/112.9x (*MSH2*), and 1.2%/98.3x (*MSH6*).

Regarding phasing, we achieved 99.88% accuracy by correctly phasing 808 out of the 809 GIAB-annotated heterozygous sites in the five loci without singularities: 308 out of 308 (*BRCA1*); 71 out of 71 (*BRCA2*); 110 out of 110 (*APC*); 195 out of 195 (*MSH2*); and 124 out of 125 (*MSH6*). The only inconsistency with GIAB data was a flip error in the *MSH6* locus—a swap of maternal and paternal alleles. Additionally, we were able to infer a single phase block with all the expected phased positions for these five ∼200 kb targets (a phase block is a contiguous set of phased locations). Four additional GIAB-annotated heterozygous sites were not recovered with sufficient reads in the *APC* and *MSH6* targets, potentially representing deletions in our batch of B cells. We did not include these four sites without coverage in our calculations. Additionally, twenty-four GIAB-annotated-but-unphased heterozygous sites were *de novo* phased in our tests: 8 sites in *APC*, 13 sites in *MSH2*, and 3 sites in *MSH6*.

For the *MLH1* target, one of the targets with singularities, genotyping was 100% accurate (16 out of 16 correctly recalled heterozygous sites), but phasing was largely incomplete with only 9 correctly phased positions (56%). We attributed this observation to the distinctively low heterozygosity of this target combined with its relatively low coverage, as mentioned above. Furthermore, we inferred a discontinuous phase block in the *MLH1* target (**Suppl. Fig. 2D**, Hap1 and Hap2 columns). We attributed these results to an insufficiency of linked reads among heterozygous sites located at least ∼10 kb apart (**Suppl. Fig. 2E**, phased sites represented by green columns, unphased sites represented by black columns). For the second target with a singularity, *PMS2*, all GIAB-annotated heterozygous sites were called as homozygous in our data, consistent with a haplotype dropout (**Suppl. Fig. 2F**). In agreement, we observed a high coverage imbalance between the two sets of GIAB-annotated alleles with an average of 91.98% and 8.02% relative abundance between the two expected haplotypes, compared to 48.52% and 51.48% in HG002 GIAB WGS data (**Suppl. Fig. 2G**). Interestingly, the alleles of the lost haplotype were still detectable to some degree (8.02%), suggesting that there is some level of inversion mosaicism affecting primarily one of the two haplotypes (Hap2 in **Suppl. Fig. 2F**), as also indicated by a multiplicity of breakpoints inferred from WGS linked-read data (**Suppl. Fig. 2C**, large heatmap).

Together, these analyses demonstrate that the targeted TELL-Seq protocol is an efficient method for phasing long, partially enriched targets with a collective accuracy close to 99.9% for five typical targets. We concluded that atypical targets, such as those with an unusually low density of heterozygous sites, would require higher relative coverage compared to typical targets to phase distally located sites and generate a complete phase block. Additionally, our analyses highlight the risk of using size-based selection as DNA pre-enrichment method when a large genomic rearrangement is not anticipated. We suspect, however, that the low resolution of pulse-field-electrophoresis (used in the HLS-CATCH system) likely enables the isolation of most inversion polymorphisms, generally smaller than a few kilobases in the human population ^31^.

### Phasing partially enriched, mid-size targets

<u>Ca</u>s9-based <u>Ba</u>ck<u>g</u>round <u>E</u>limination (CaBagE) is a non-size-based DNA-pre-enrichment method that leverages the high stability of Cas9/gRNA bound to DNA post cleavage to protect the two ends of the target from subsequent exonuclease digestion ^33,34^. Meanwhile, (unprotected) non-target DNA is readily digested ^33,34^. Using a modified CaBagE version, we sought to enrich and phase four additional targets. Since we used a salting-out method to extract genomic DNA that includes pipetting and centrifugation steps (see Methods) ^35^, we expected to obtain DNA with shorter HMW properties than those observed with gently extracted genomic DNA with the HLS-CATCH system. Therefore, we focused on relatively small target fragments (∼20 kb), which we designed to be adjacently located along the *MSH2* locus (hereafter referred as T1-T4). Targeting adjacent loci allowed us to use the smallest possible number of Cas9/gRNA complexes (five) to enrich the four targets simultaneously (gRNA #1-#5, **Fig. 2A**). This design required us also modifying the CaBagE approach by using a catalytically inactive Cas9 form—deactivated (d)Cas9—to avoid Cas9 from remaining stably bound to only one side (5’ or 3’) of the gRNA binding site at the T1-T2, T2-T3, and T3-T4 intersections after cleavage. Retaining dCas9 at both sides of the gRNA binding site enables the protection towards the direction with a second RNP complex at the other end of the DNA molecule (gRNA #1-#5, **Fig. 2A**) ^36^. Such gRNA distribution would also enable the recovery of ∼40 kb, ∼60 kb, and ∼80 kb fragments if fragments of these sizes were preserved during DNA extraction (**Fig. 2A**) ^37^.

**Figure 2.**
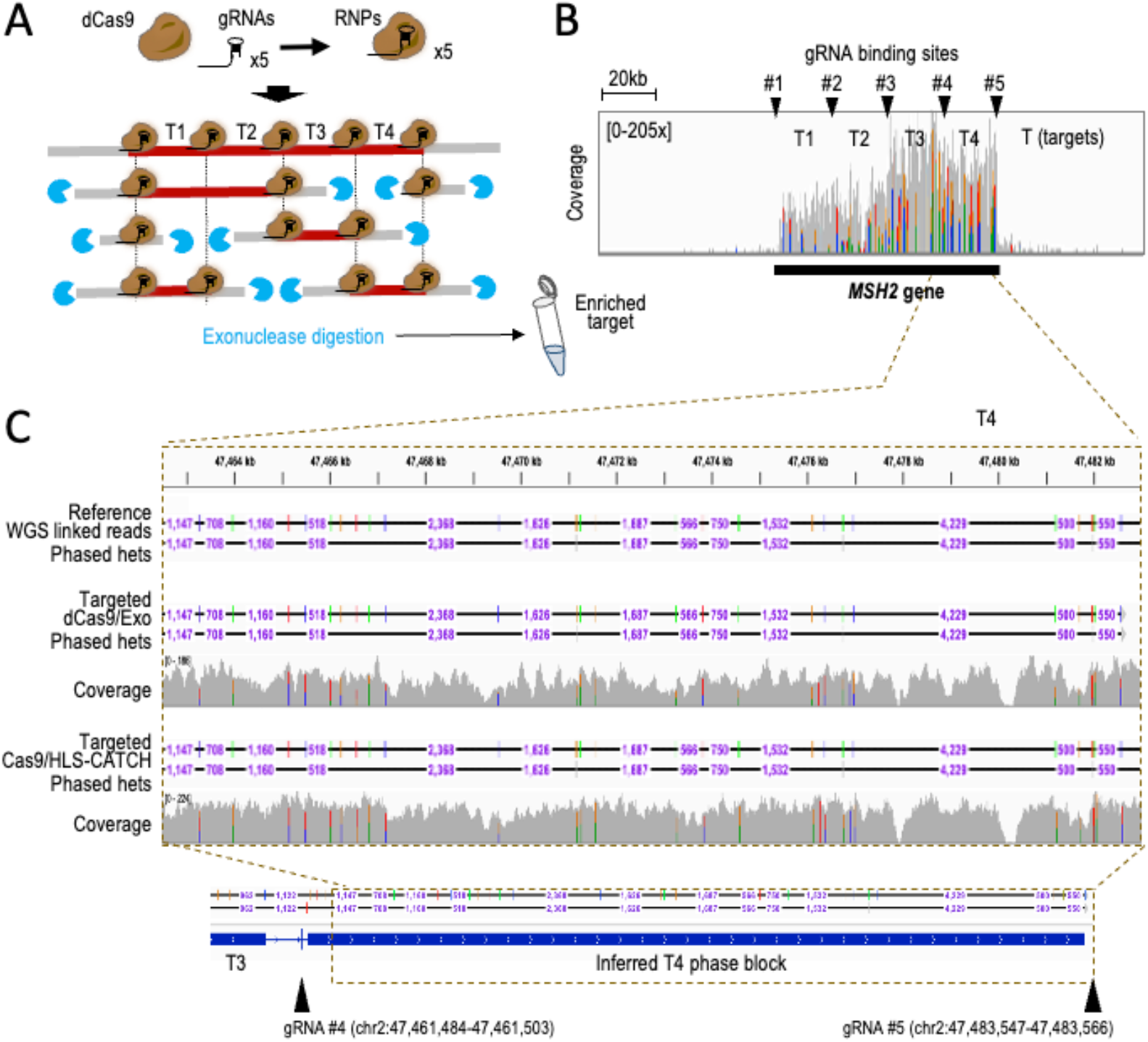
TELL-Seq-mediated phasing of four adjacent ∼20 kb targets enriched with a modified CaBagE method that leverages dCas9 to protect both sides of the targeted regions. **A**. Experimental workflow. Targets can be enriched as individual loci (∼20 kb) or as longer ∼40, ∼60, or ∼80 kb fragments, depending on the HMW properties of input genomic DNA. Protected DNA (target) is represented in dark red; unprotected DNA (non-target) is represented in grey. **B**. Coverage across the *MSH2* locus (black bar indicates *MSH2* gene position). This profile supports high on-target recovery over background for all targets (T1-T4) but higher for T3-T4 than for T1-T2. Arrowheads indicate gRNA binding sites (#1-#5). **C**. Screenshot from the IGV Portal showing phased data and coverage using Targeted TELL-Seq with dCas9/exonuclease or HLS-CATCH/Cas9 as DNA pre-enrichment methods. Shown on top HG002 WGS linked-read phased data (benchmark). Phased data includes distances between phased sites (we note that distance are not indicated when not allowed at the selected resolution on IGV). Bottom tracks show phased data and inferred phase blocks with Targeted TELL-Seq using modified CaBagE. Arrowheads indicate gRNA binding sites.

After RNP assembly, we incubated the pool of five RNP complexes (#1-5) with 1 μg of HG002 HMW genomic DNA followed by exonuclease digestion. We then applied the TELL-Seq protocol used for HLS-CATCH-enriched targets to approximately half of the enriched DNA material. After sequencing and duplicate removal, we obtained 6.3 million reads with 0.8% on-target recovery and 68.7x average coverage depth.

Among the four targets, T3 and T4 were more efficiently recovered than T1 and T2 (0.3%/104.5x and 0.3%/85.6x, compared to 0.1%/40.5x and 0.1%/44.9x, respectively), suggesting that RNPs #3-5 provided better protection than RNPs #1-2 (**Fig. 2B**). Further examination of the data revealed 100% recall and phasing accuracies in T3 and T4: 27 out of 27 and 32 out of 32, respectively (**Fig. 2C**, T4). Two independent phase blocks were inferred in T3 and T4, which suggested that ∼40 kb and larger fragments were rare or inefficiently processed in the sample (**Fig. 2C**, bottom track). For T2, three GIAB-annotated heterozygous sites (out of 33) were not covered by any read, potentially due to a short deletion in our batch of cells (**Suppl. Fig. 3A**). The rest of heterozygous sites in T2 (30) were correctly recalled and phased except for one site that remained unphased (**Suppl. Fig. 3B**); together, representing a ∼96.7% phasing accuracy (29 out of 30). For T1, phasing was more problematic. Despite that all heterozygous sites (9 out of 9) were recalled, two sites remained unphased, leading to a ∼77.8% phasing accuracy (**Suppl. Figs. 3C** and **D**). Additionally, we inferred a discontinuous phase block for T1 (**Suppl. Figs. 3C** and **D**, bottom track). We attributed these problems to a distinctively low heterozygosity in this target (9 heterozygous sites in T1 compared to 27, 32, and 30 in the other three targets of the same size, T2-T4).

Together, we correctly phased 95 out of the 98 recalled heterozygous positions, which represents a ∼96.9% phasing accuracy. The unphased sites were associated with a low heterozygosity density combined with a relatively low coverage of the T1 target. This demonstrates that TELL-Seq is capable of accurately phasing partially enriched targets isolated with a Cas9-based DNA enrichment method that does not rely on size for target isolation. Moreover, our analyses of the T1-T4 targets did not result in any flip or switch phasing errors.

### Minimizing DNA collisions with highly pure targets

Next, we sought to validate relatively short, highly pure targets (e.g., amplicons) as input material for targeted TELL-Seq. Phasing amplicons will require further optimization of the targeted TELL-Seq protocol. The expected high purity of an amplicon is incompatible with conditions that allow collisions during the DNA co-barcoding reaction. Otherwise, it will be likely that maternal and paternal copies of the target collude on the same microbead. To identify collision-free but still cost-effective co-barcoding conditions (i.e., without using an excessive volume of microbeads), we processed five TELL-Seq libraries with 20, 50, 100, 200, and 400 pg of the BstP I-digested lambda phage genome. BstP I digestion splits the 48.5 kb lambda phage genome into fourteen non-overlapping fragments, which mimics a pool of 0.1-8.5 kb targets with identical number of molecules in every case (117, 224, 702, 1,264, 1,371, 1,929, 2,323, 3,675, 4,324, 4,822, 5,687, 6,369, 7,242, and 8,453 bp). This diversity should allow us to detect collisions with the caveat that we would miss situations in which a collision occurs between two molecules of the same fragment. After sequencing, we recovered all the ‘targets’ except the smallest 117 bp fragment (**Suppl. Fig. 4A**). On-target recovery correlated with fragment size (R^2^=0.8111-0.8461; **Fig. 3A**), but the 100 bp fragment ends showed distinctively lower coverage (**Suppl. Fig. 4B**). To estimate the fraction of collision-free barcoding events, we capitalized upon the sequence differences among the fourteen fragments, observing the highest collision-free fraction for 20 pg inputs (84.8±6.5%) and the lowest collision-free fraction for 400 pg inputs (38.7±4.8%), gradually decreasing with increasing DNA amounts: 50 pg, 70.7±4.9; 100 pg, 61.0±4.7; and 200 pg, 46.5±5.1 (**Fig. 3B**). Within the fraction of estimated collision-free barcoding events, we also calculated linked-read efficiencies—linked reads over total short reads. Linked-read efficiency was the highest with 20 pg inputs (40.9±10.9%) and the lowest with 400 pg inputs (29.4±11.3%) but relatively uniform between both amounts: 50 pg, 40.6±11.2%; 100 pg, 37.7±9.6%; and 200 pg, 35.4±11.4% (**Fig. 3C**). As expected, linked-read efficiencies increased with coverage (as shown in **Suppl. Fig. 4C** after subsampling all sequencing outputs at 75%, 50%, 25%, and 10%). In general, linked-read efficiencies showed no strong dependency on fragment size when normalized by coverage for 0.7 kb and larger fragments (**Suppl. Fig. 4D**).

**Figure 3.**
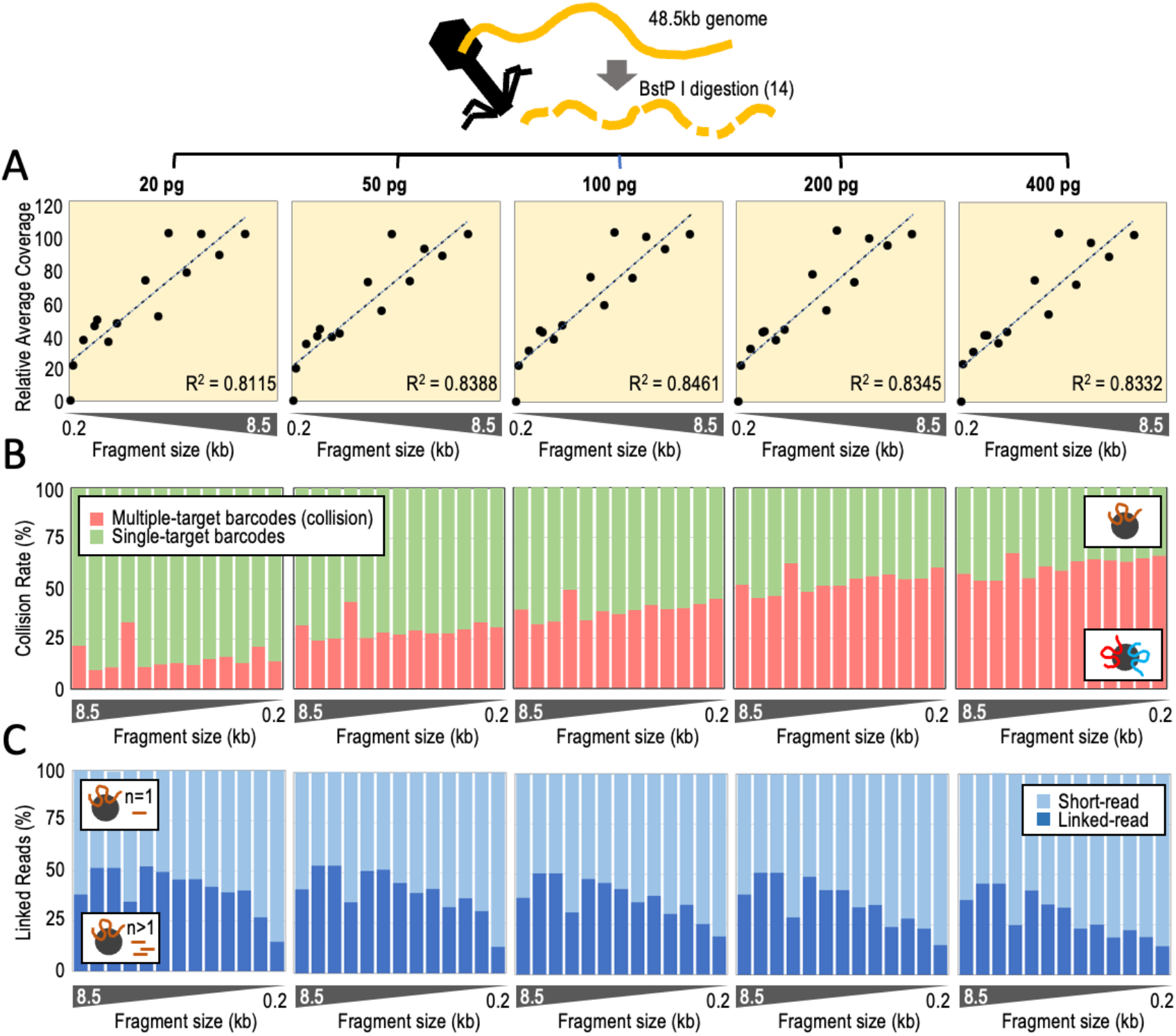
Determining DNA collision rates and linked-read efficiencies using BstP I-digested lambda phage genomic DNA. **A**. Relative coverage (average across every fragment) relative to fragment size. On top indicated input amounts of digested fragments. Y-axis represents the average coverage relative to the average coverage with the 8.5 kb fragment, X-axis represents fragment size (each datapoint is one fragment). **B**. DNA collision rates (%) relative to fragment size by input amounts. **C**. Percentage of linked reads relative to the total number of short reads by fragment size and input amounts. Analysis limited to microbeads with only one associated fragment (single-barcode fragments or collision-free cases).

Together, these analyses suggest that using 20 pg target inputs allows ∼85% collision-free DNA co-barcoding events. The caveat is that, in our experience, inputs of 20 pg and lower can generate inconsistent library yields. We, therefore, reasoned that consistency in library yields could be achieved by adding 60-80 pg non-human DNA to 20 pg target inputs, hereafter referred to as ‘filling’ DNA (e.g., *Escherichia coli* or BstP I-digested lambda phage genomic DNA). Choosing these conditions would be a compromise between minimizing DNA collisions and maximizing library-yield consistency and cost-effectiveness (with the lowest possible number of microbeads but also having enough molecules of the target). When there are four or more targets in a sample for library preparation, filling DNA is not required.

### Phasing amplicons

To test whether 20 pg inputs supplemented with 60-80 pg of filling DNA are conditions for phasing, we generated a set of long PCR products to use as input material in targeted TELL-Seq reactions. Long-range PCR (3-30 kb) can be a notoriously challenging process, especially when amplicons are ultimately used for the purpose of phasing. First, long-range PCR often requires substantial optimization using additives and a polymerase with enhanced DNA binding properties and the highest fidelity and processivity ^38,39^. Second, PCR conditions must be established that reduce the risk of haplotype dropouts and chimeras, both problematic for phasing. Haplotype dropouts can be generated when either the maternal or paternal copy of the target is inefficiently amplified ^40^. Chimeras—a crossover of maternal and paternal haplotypes—can occur when an incomplete PCR product acts as a primer on the wrong amplicon copy during PCR amplification, leading to a switch error—an artificial haplotype ^41^. In part, the risk of haplotype dropouts and chimeras can be reduced by using long extension times at the end of every PCR cycle and by avoiding over-amplification. In this context, it is advantageous that TELL-Seq requires only picogram amounts of input DNA, which allows minimizing the number of PCR cycles.

With these precautions, we amplified a 13 kb region in the *SCN5A*-*SCN10A* locus containing seven polymorphic sites near gene-regulatory elements associated with a high risk for sudden cardiac arrest and a well-characterized haplotype diversity with at least fifteen reported allelic combinations (Hap1-15) ^42–44^. These seven sites are spaced 288, 4,299, 23, 69, 5,560, and 2,505 bp apart (from 5’ to 3’) ^44^. As a second target, we selected the 3.4 kb region spanning the exons of the *PIK3CA* gene. *PIK3CA* is the second most frequently mutated gene across all cancer types, being mutated at least twice in 8-13% of clinical cases ^45,46^. Phasing double *PIK3CA* mutations can help to predict oncogenicity and sensitivity to treatment, such as when the E545K and L866F mutations coincide along the same chromosomal copy (in cis) ^46^. These mutations can be in a different chromosomal copy (in trans) relative to a nearby allelic variant, I391M (rs2230461) ^46^. The distances between these three sites are 459 and 964 bp in cDNA. We also generated shorter PCR products within the 13 kb and 3.4 kb regions to examine the reproducibility of the phasing data: 3.9, 4.1, and 2.7 kb for *SCN10A* (from 5’ to 3’); and 1.8 kb for *PIK3CA* (**Fig. 4A**).

**Figure 4.**
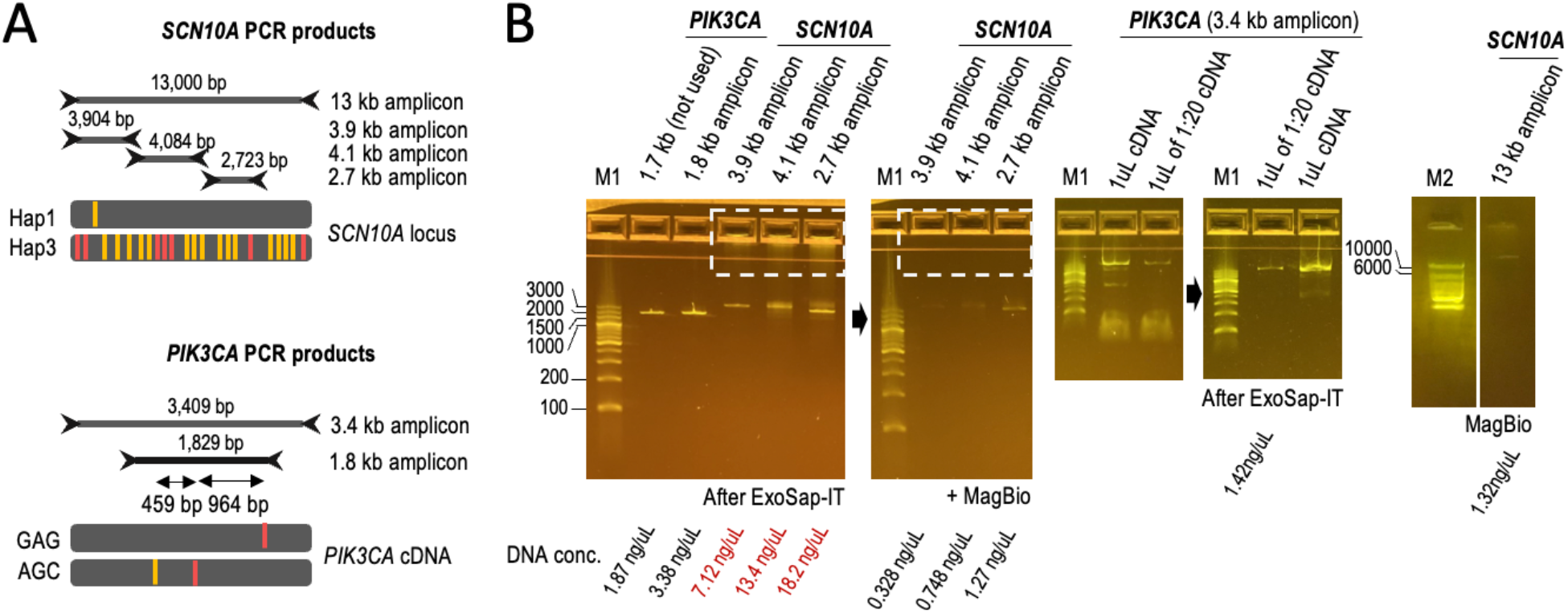
Amplicons used in this study. **A**. *SCN10A* and *PIK3CA* amplicon sets phased in this study (13, 3.9, 4.1, and 2.7 kb from genomic DNA and 3.4 and 1.8 kb from cDNA, respectively). HG001 carries previously annotated Hapl and Hap3 haplotypes. Red bars indicate the position of 7 clinically relevant heterozygous sites; orange bars indicate the position of 15 additional heterozygous sites. Alleles are colored if represent the alternate allele, i.e., different from the human reference. HCC202 cells carry two clinically relevant alleles (in red) and one non-clinically relevant variant (in orange). **B**. E-gels with PCR products as indicated. Non-specific DNA and primers were removed as indicated (see dashed square as example). We selected minimally amplified PCR products (see PKI3CA example with 1:20 diluted cDNA). DNA concentrations are indicated at the bottom.

We used the GIAB reference HG001/NA12878 as a source of genomic DNA to amplify the subset of *SCN10A* targets. WGS linked-read data ^24,32^ reveals that HG001 is a Hap1/Hap3 carrier in the *SCN5A*-*SCN10A* locus (TACCATT and CGGGGGC) ^44^. WGS linked-read data ^24,32^ also reveals that HG001 carries another subset of fifteen heterozygous sites, some of which have also been recently associated with heart arrhythmia ^42^. We have observed that the twenty-two heterozygous sites are split by major and minor allele frequencies, with all major alleles in one chromosomal copy and all minor alleles in the other chromosomal copy. This finding builds upon the previously proposed model that the 13 kb Hap1/Hap3 region represents Ying-Yang haplotypes ^44^. To generate *SCN10A* amplicons, genomic DNA was extracted with the same modified salting-out method used to isolate the ∼20 kb T1-T4 targets (**Fig. 2A**). To generate *PKI3CA* amplicons, we used cDNA generated from human breast cancer HCC202 cells. HCC202 cells carry the GAG and AGC haplotypes underlying the three heterozygous sites under investigation ^46^. All amplicons appeared as weak, mainly single PCR products on E-gels (before cleanup); thus, indicating no over-amplification (**Fig. 4B**). Additionally, amplicons were generated without major haplotype biases in most cases, including for 13 kb PCR products (**Suppl. Fig. 5**).

We generated forty TELL-Seq libraries to assess different amplicon amounts, target sizes, and cDNA priming methods, as well as to acquire replicates (between 2 and 6). We generated thirty-four libraries with a single amplicon, and six libraries with a panel of amplicons (3.4 kb *PIK3CA* and 2.7, 3.9, and 4.1 kb *SCN10A*). The input amounts varied from 20 pg to 5 pg and 0.04 pg and were supplemented with filling DNA when processing single targets (either *E. coli* genomic DNA or BstP I-digested lambda phage genomic DNA). For target panels (4 amplicons), we used equal amounts for each target (20 pg) without supplementing with filling DNA. The rationale behind using equal amounts for each target (rather than equal molecule numbers) is that—we postulated— low coverage for the smallest fragments would be compensated with a higher molecule number compared to larger fragments. We tested cDNA-based products with reverse transcriptase primed by random hexamers and oligo-dT. We mapped *SCN10A* reads to genomic DNA and *PIK3CA* reads to *PIK3CA* cDNA with the allele frequency (AF) cutoff set to 0.1 (see Methods).

Analysis of the sequencing data revealed that the average on-target coverage ranged between 58.8x and 3,574x with most experiments falling between 200x and 800x (**Fig. 5A**, average on-target coverage). Out of the 366 annotated heterozygous sites—collecting the numbers from the fifty-eight replicates—we correctly phased 364, while the remaining two sites represented flip errors, overall achieving 100% and 99.45% genotyping and phasing accuracy, respectively (**Figs. 5A** and **5B**). Importantly, the two flip errors were not reproduced in replicate samples. Additionally, we identified (and phased) a heterozygous site not previously annotated by GIAB, which was consistent across fifteen libraries and may, therefore, represent a genuine variant specific of our cells (**Fig. 5A**, *de novo* phasing). Together, these results demonstrate that targeted TELL-Seq can achieve highly accurate phasing of amplicons and can even reach a perfect accuracy when results are supported by replicates.

**Figure 5.**
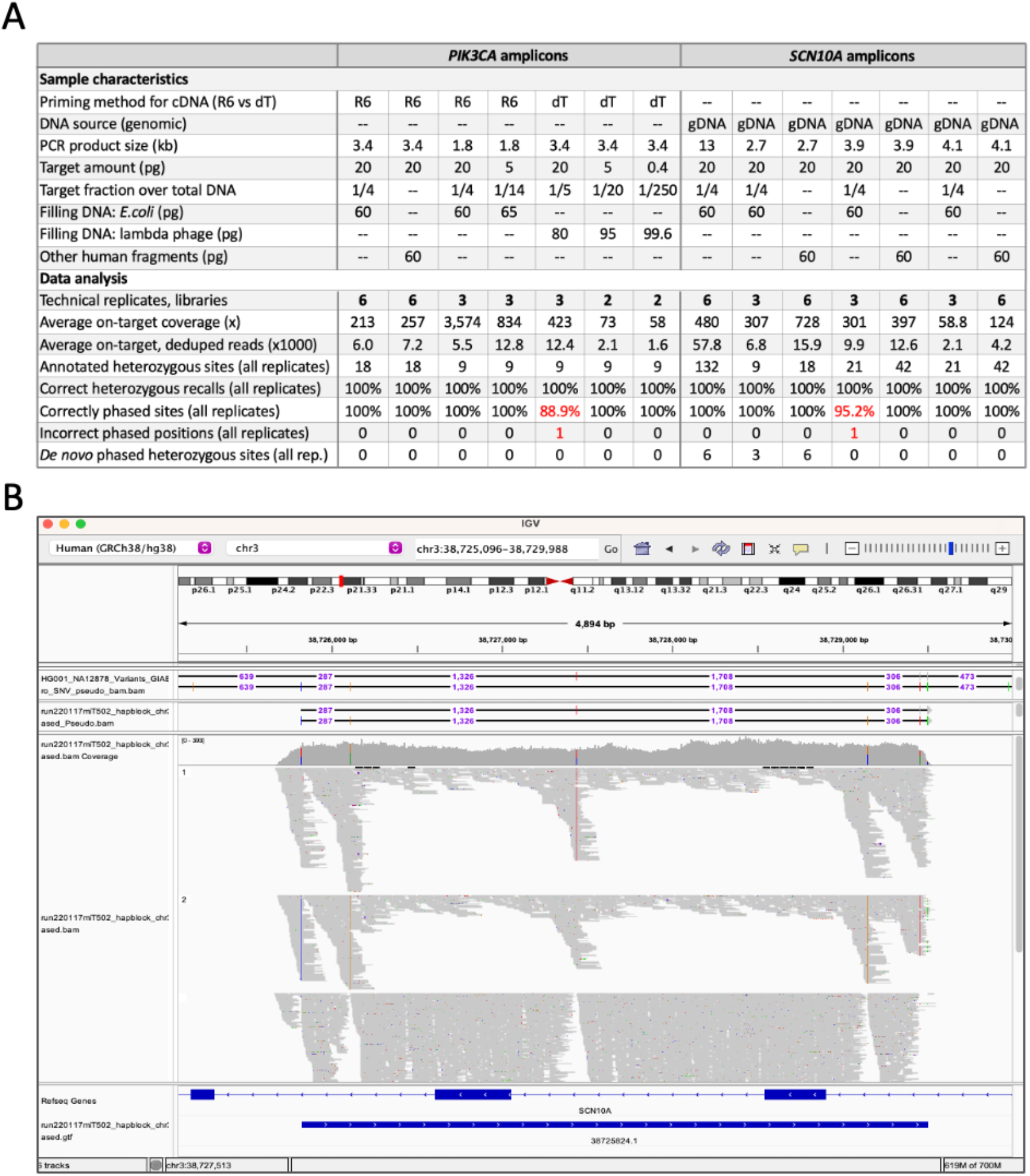
Phasing *SCN10A* and *PIK3CA* amplicons (sizes: 1.8-13 kb). **A**. Summary table of genotyping and phasing results using *SCN10A* and *PIK3CA* amplicons as input DNA; PCR from NA12878 genomic DNA (n = 33 tests) or breast cancer HCC202 cDNA (n = 25 tests). In red, highlighted the phasing errors. R6 (random hexamers) and dT (oligo-dT) for cDNA priming, as indicated. B. Screenshot showing the phasing of three technical replicates on the IGV Portal. Fragments corresponds to 4.1 kb *SCN10A* amplicon. The top track shows the phasing of one 4.1 kb replicate in phased .pseudo .bam format. Heterozygous sites are separated by numbers that represent distances between neighboring sites. The third track show coverage, inferred haplotype 1 reads, inferred haplotype 2 reads, and unphased reads. The last tracks shows gene annotations and an inferred phase block (blue bar).

We next benchmarked targeted TELL-Seq with previously generated ONT data of 13 kb *SCN10A* products amplified from peripheral blood-derived genomic DNA from three Hap1/Hap3 carriers (referred here to as individuals #1-3) ^44^. Our results agreed with the previously reported genotypes in individuals #1 and #2, correctly recalling and phasing the 14 (7+7) clinically relevant heterozygous positions in these samples (**Fig. 6A**, Ind.#1 and #2). For individual #3, the seven sites were also recalled as heterozygous in our data, although the most 5’ site remained as unphased and the most 3’ site was phased inconsistently with prior data (**Fig. 6A**, Ind.#3, Replicate #1). Since coverage was lower for individual #3 than individuals #1-2 (54.6x compared to 89.9x and 92.0x, respectively), we suspected that our call for the most 3’ site in individual #3 was erroneous. In agreement, we generated a second TELL-Seq library from the same amplicon (Ind. #3, Replicate #2) and reamplified the first TELL-Seq library from the remaining target-bound microbeads (Ind. #3, Replicate #1.2). These two new analyses agreed with a Hap1/Hap3 genotype and were based on higher coverage (105.1x and 290.4x; **Fig. 6A**, Ind.#3, Replicate #2 and #1.2).

**Figure 6.**
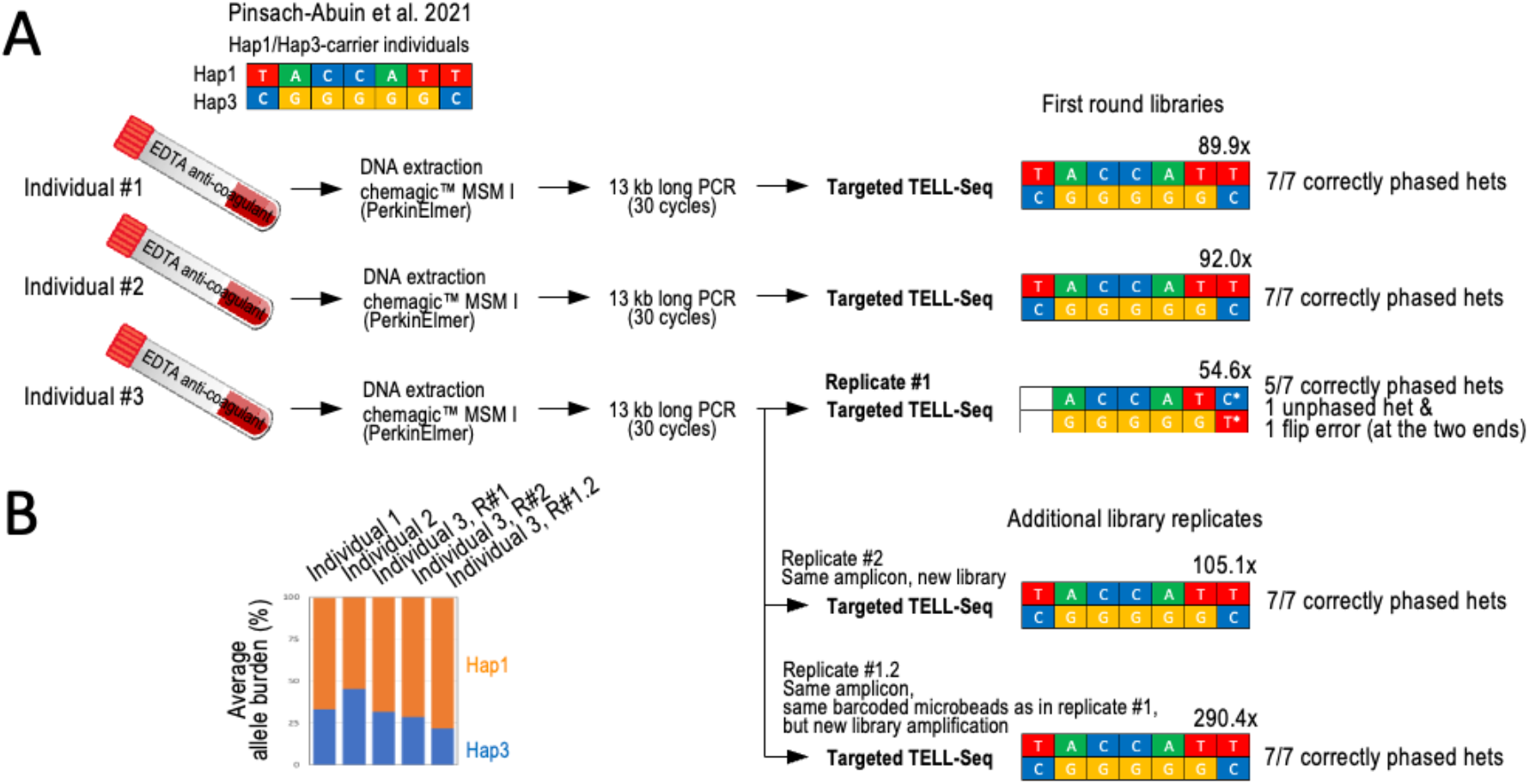
Targeted TELL-Seq with three 13 kb amplicons generated from peripheral blood-extracted genomic DNA from three known Hap1/Hap3 carrier individuals. **A**. Previous work identified the three Hap1/Hap3 carrier individuals from whom 13 kb amplicons were generated and now have been processed in this study. We confirmed the Hap1/Hap3 genotype in all three individuals with Targeted TELL-Seq. However, this conclusion needed the three replicates for Individual #3. In Replicate #1, we observed an unphased and a flip error. We note that this library provided the lowest coverage, 54.6x (Individual #3). We generated a new library from the same amplicon (Replicate #2) and re-do the library amplification with leftover barcoded microbeads from Replicate #1 (Replicate #1.2). In both cases, we validated the expected Hap1/Hap3 genotype. We note that the sequencing depth was also higher for both replicates, 105.1x and 290.4x, respectively. Coverage indicated (x) on top of each genotype and number of correctly re-called and phased heterozygous positions in Hap1 and Hap3 are also indicated. **B**. Graph shows average allele burden for all Hap1/Hap3 sites by library. In all cases, Hap1 was amplified slightly more efficiently than Hap3, although it had no effect on genotyping and phasing.

We noticed that linked-read coverage at the two haplotype ends in the first replicate from individual #3 had been the lowest of all tests, which indicates again a connection between phasing problems and low coverage (**Suppl. Fig. 6A**, orange columns). For reference, ONT coverage was at least ∼600x for long reads containing all seven clinically relevant heterozygous sites (**Suppl. Fig. 6B**) ^44^. Intriguingly, we observed a slight haplotype bias in our tests with blood-derived samples (Hap1>Hap3), which we did not observe when amplifying the 13 kb fragment from HG001 (compare allele burden plots between **Suppl. Fig. 5 and Fig. 6B**). However, this haplotype bias did not apparently have a negative impact on the process of phasing.

### Minimum sequencing depth for accurate phasing

Finally, we sought to examine how sequencing coverage (used as a proxy of linked reads) affects genotyping and phasing accuracy when following the targeted TELL-Seq protocol. To do this, we subsampled nine of the sequencing outputs generated from the libraries using amplicons as input material and re-did the phasing analyses (outputs subsampling at 50%, 25%, 12.5%, 6.25%, 3.125%, 1.56%, and 0.78% of the original dataset). After duplicate removal, we segregated the data into four groups based on phasing quality. Group 1 represents data without genotyping or phasing errors and a single phase block; group 2 represents data without genotyping errors and a single phase block but with one flip or switch error; group 3 represents data with incorrect genotyping, discontinuous phase blocks, and flip/switch errors; and group 4 represents data with mostly unphased heterozygous sites (**Suppl. Fig. 7**). Grouping the subsampled data in this way revealed that lowering the coverage below 180x (for the 13 kb fragment) and 150x (for the 2.7-4.1 kb fragments) led to phasing errors and that further lowering the coverage led to genotyping errors (**Suppl. Fig. 7**, compare group 1 and groups 2-4). However, we note that data at 50-60x coverage may often show no phasing errors, as we have frequently observed (**Fig. 5A**). Together, these subsampling analyses highlight the importance of sufficient coverage to reduce the risk of phasing errors.

## DISCUSSION

Long-range information embedded into short reads is essential for interpreting genomic data for many different applications ^17,47–54^. Single-tube TELL-Seq is an instrument-free linked-read method characterized by its versatility and simplicity ^24^. It leverages a unique transposition chemistry to enable millions of simultaneous long-range, information-capturing reactions on the surface of micron-sized beads ^24^. The WGS TELL-Seq protocol is highly permissive of DNA collisions (up to 6-8 HMW DNA fragment, on average, when processing a human genome). It is highly unlikely that a maternal and paternal copy of the same locus will coincide on the same microbead when processing an entire human genome. However, when phasing an enriched locus, TELL-Seq must minimize DNA collisions due to the low complexity of the sample. In this study, we present targeted TELL-Seq for cost-effective phasing of discrete DNA fragments ranging in sizes between 1.8 kb and 200 kb. This protocol has a version for impure targets (<1-2%; e.g., when enriched with Cas9-based methods) and a version for pure targets (>90-95%; e.g., with amplicons). Both versions can be used with single targets or target panels with only minor procedural variations, as here described. Pooling targets (panels) is the most cost-effective phasing strategy. However, it is important to highlight that the target sequences must not overlap in the reference genome.

We benchmarked targeted TELL-Seq using WGS linked-read and amplicon-based long-read data, demonstrating high-quality genotyping and phasing accuracies. For example, with impure ultra-long targets, we correctly phased 808 out of 809 heterozygous sites after accumulating data from five ∼180-200 kb targets without singularities, a phasing accuracy of 99.88%. In a case with an abnormally low heterozygosity, we observed that the sequencing depth that can phase a target with a typical heterozygosity density, it can only phase the subset of closely separated heterozygous sites (<10 kb). For another case with an ultra-large inversion, phasing is problematic when using a size-based DNA-pre-enrichment method. As a solution to these problems, we recommend sequencing deeper and redesigning gRNAs, generating genome-scale TELL-Seq data, or—if the target is relatively short—using an alternative, non-size-based target pre-enrichment method, respectively. With mid-sized targets, we achieved a 98.88% phasing accuracy (88 out of 89 heterozygous sites correctly phased) with three ∼20 kb targets. In a fourth target with low heterozygosity compared to the other three phased loci, however, we detected unphased sites and a discontinuous phase block. With 2-13 kb amplicons, we achieved a phasing accuracy of 99.45%. (364 out of 366 heterozygous sites correctly phased). We also correctly phased 28 out of 28 clinically relevant genomic sites with blood-derived PCR products, which is a 100% phasing accuracy, although we did not include in this calculation a case that had phasing issues due to a relatively low sequencing depth. In summary, we have always generated robust data unless the sequencing depth was insufficient due to low heterozygosity density. Most phasing issues were noticeable by unphased heterozygous positions and discontinuous phase blocks, which should prompt for generating a new library and sequencing deeper. Of note, erroneous phasing was rare and non-reproducible, which provides a solution to bypassing this problem: make conclusions based on reproducible results.

Since sequencing depth is critical to phase a target, we wondered how much on-target sequencing is required to correctly phase a target. In general, our data suggest that it will depend on heterozygosity density, but the risk of phasing errors or incomplete phasing is the lowest above 150x of on-target coverage. However, we have also found many examples with coverage lower than 150x and correct and complete phasing. We would recommend, therefore, generating replicate libraries to compare results and reach to conclusions. With amplicons, PCR conditions would also be important. The risk of chimeras and allelic dropouts can be reduced by avoiding over-amplification. Replicates could also be generated using different sets of primers, as we show with the *PKI3CA* target. Additionally, multi-band PCR products should be avoided unless there is confidence that the extra products do not represent shorter versions of the target. Notably, no major amplification biases were observed in our tests, except for blood-derived PCR products, but the bias appeared not to confound the phasing results.

Regarding input amounts, we have established that high-quality phasing data can be obtained with no more than 100 pg of an impure target and 20 pg of a pure target. When working with <80 pg inputs, we recommend supplementing with filling DNA for yield consistency and the highest recovery. The lowest target input amount tested is 0.08 pg (supplemented with 99 pg of filling DNA), suggesting that, at least in principle, hundreds of (non-overlapping) targets could be pooled in a single TELL-Seq library preparation (especially valuable for Cas9-based DNA pre-enrichment methods using hundreds of gRNAs). Filling DNA should be either bacterial genomic DNA or digested lambda phage DNA split into at least 10-14 fragments (regardless of restriction enzyme/s used; we used BstP I, but other options could be used generating at least 10-14 fragments).

Regarding the DNA-pre-enrichment method, our data confirm the value of using a gentle, pipetting-free system, such as HLS-CATCH, that preserves ultra HMW DNA properties and isolating targets in pools. The caveat of using a system that relies on size for target isolation is that unexpected genomic rearrangements that significantly alter the size properties of the target may lead to haplotype dropouts. Nonetheless, most genomic rearrangements should not have a major effect on DNA motility in pulse-field electrophoresis, since it is a low-resolution method.

Finally, a note about the IGV tracks showing phased TELL-Seq data throughout this study (e.g., **Figs. 1D, 2C**, and **4B**). These tracks were generated using a new computational tool that we have recently developed that allows convenient browsing of phased variants as part of haplotypes and phase blocks and enables to examine the distance between heterozygous sites and match data with gene annotations and read coverage on the IGV Portal. This tool is now available through the Tell-Sort pipeline for TELL-Seq analysis (see Methods).

Targeted TELL-Seq represents a substantial gain in cost-effectiveness and analysis simplicity when phasing a target. For reference, we observed similar phasing results with 20 million reads generated by targeted TELL-Seq as with 2 billion reads generated by WGS TELL-Seq or WGS 10x Genomics linked-reads, which is a hundred times less sequencing burden for the targeted option. In addition, the difference in computation time between the targeted and WGS options is remarkable, taking only a few minutes for data generated with a targeted approach compared to a few days for a genome-scale analysis. In conclusion, TELL-Seq empowers researchers working with the most popular sequencing platforms (short-read-based) to phase targets in the 2-200 kb range and likely shorter and longer without the need of a long-read platform.

## Limitations of the study

While our results indicate that TELL-Seq is an efficient method for phasing targets with extremely low error rates, it remains to be determined whether the same performance is observed with other targets and DNA enrichment methods and amplification conditions not tested in this study. Additionally, it should be noted that TELL-Seq is solely intended for research use and not for use in clinical applications or diagnostic procedures despite our validation tests were conducted with clinically relevant targets.

## ACKNOWLEDGMENTS

We thank Diogo Comprido from NZYTech (Portugal) for providing NZY reagents and Professor Connie C. Bezzina from Amsterdam University Medical Centers (Netherlands) for providing association data from individuals diagnosed with heart arrythmia. This work was funded by Universal Sequencing Technology Corp.

## Author contributions

Conceptualization, Z.C. and I.G.B.; investigation, V.M., M.R., T.K.; reagent generation, R.Z., L.P., L.Y., A.P.A., S.P., and C.B.; data analysis, I.G.B., Y.X., and P.L.C.; algorithm development, Y.X, P.L.C., and Y.W.; writing – original draft, I.G.B; writing – review & editing, I.G.B., Z.C., V.M., M.R., P.L.C., Y.X., T.K., and N.M.; funding acquisition, M.L. and Z.C.

## MATERIALS AND METHODS

### Human cell sources

Human lymphoblastoid GM24149, GM27730, and GM12878 cells were purchased from Coriell Cell Repositories^®^ (Cat. # GM24149, # GM27730, and #GM12878). Human epithelial breast cancer HCC202 cells were purchased from ATCC^®^ (Cat. #CRL-2316).

### Peripheral blood-derived amplicons from human donors

Amplicons from three unrelated individuals were previously generated from peripheral blood-derived genomic DNA ^44^. Briefly, blood was collected in EDTA anti-coagulant BD Vacutainer tubes and genomic DNA was extracted using the Chemagic MSM I Instrument (PerkinElmer) following the manufacturer’s recommendations and stored at −20°C before PCR ^44^. The samples were obtained according to the Declaration of Helsinki Principles and complying with the European and National Code of Practice with consents approved by the Clinical Research Ethics Committee of the Dr. Josep Trueta Hospital (#2012.097).

### Genomic HMW DNA extraction and target isolation with the HLS-CATCH system

For HMW genomic DNA extraction and target isolation with the HLS-CATCH system (Sage Sciences), we followed instructions according to the manufacturer’s recommendations ^27^. We used one million cells for each isolation. We performed three isolations: *BRCA1* and *BRCA2* separately and *APC, MLH1, MSH2, MSH6, PSM2* collectively using GM24149/HG003 cells for *BRCA1* and *BRCA2* and GM27730/HG002 cells for the rest of the targets. Cas9/gRNA RNP complexes were assembled with 2μM Cas9 and the pool of gRNAs (see table below). After 4 min/80 V electrophoretic injection, each targeted fragment was size separated and eluted by pulse-field electrophoresis. This approach yielded 220,000-400,000 copies per targeted locus with an enrichment of 200-400-fold over genomic DNA (measured by quantitative PCR). The isolated DNA fractions ranged between 3 and 7 ng.

### HMW genomic DNA extraction with a modified salting-out method

For GM12878/HG001/NA12878 and GM27730/HG002/NA24385, HMW genomic DNA was extracted using a salting out method previously reported ^35^ with some modifications. Five million cells were harvested at 315xg for 7 minutes in a 15 mL centrifuge tube (Beckman Coulter Allegra X-14R) and stored at -80°C. The cell pellet was resuspended in 3 mL of nuclei lysis buffer (10 mM Tris-HCl, 400 mM NaCl, and 2 mM EDTA, pH 8.0) by gentle inversion (20 times). For cell lysis and protein digestion, 0.2 mL of 10% SDS and 0.5 mL of Proteinase K solution (1 mg/mL Proteinase K, 2 mM EDTA, pH 8.0) were added to resuspended cells and mixed by gentle inversion (5 times), then incubated with rotation at 37°C overnight (12-18 hours). For DNA extraction, 1.2 mL of 5 M NaCl were first added and mixed for 15 seconds by inversion, cellular/protein debris were then collected by centrifugation at 1,000xg for 15 minutes at 4°C (Beckman Coulter Allegra X-14R), and the supernatant (DNA) was finally transferred slowly to a new 15 mL tube using a serological pipette. For DNA precipitation, 8 mL of 100% ethanol were added (200 proof [absolute] for molecular biology) and mixed by inversion (at least 10 times). A precipitate of long DNA strands should be clearly visible during this step. The DNA solution was pelleted by centrifugation at 6,000xg for 5 minutes at 4°C (Eppendorf Centrifuge 5425R). After removing the supernatant, DNA pellets were left to dry for 5 minutes at room temperature and resuspended in 300 μL of 0.1x TE buffer (10 mM Tris-HCl pH 8.0, 0.1 mM EDTA). Following an overnight incubation at room temperature and a 2-day incubation at 4°C, the DNA concentration was determined to be 50 ng/μL by the Qubit High-sensitivity dsDNA Assay (Thermo Fisher Scientific). HMW genomic DNA was stored at 4°C for up to 2 weeks or at -20°C for up to 6 months. We note that the main difference between the method used here and previously published ^35^ is in the DNA extraction step, which added pipetting and centrifugation steps after ethanol addition. If ultra-long HMW DNA is required, we suggest following the original protocol, which avoids mechanical DNA shearing by physically transferring the precipitated DNA strands with a plastic spatula or pipette directly into 200 μL of 0.1x TE buffer, yet this option has the risk of losing DNA.

### Target excision and isolation with a modified CaBagE method

We modified the CaBagE method ^33,34^ and designed a tiling strategy of spacing gRNAs as previously shown ^37^. Five gRNAs were designed to isolate four ∼20 kb adjacent targets (T1-T4) from the human *MSH2* locus. We designed gRNAs using the online tool CRISPR-Cas9 Guide RNA Design Checker provided by the Independent DNA Technologies (IDT) (https://www.idtdna.com/site/order/designtool/index/CRISPR_SEQUENCE). We pasted 0.5-1 kb DNA regions where we sought to identify gRNA binding sites and selected 20-bp sequences immediately preceding the protospacer-adjacent motif (PAM) for *Streptococcus pyogenes* Cas9 (5’-NGG-3’) without repetitive elements (e.g., Alu elements) or high-frequency single nucleotide polymorphisms (based on dbSNP13 entries). Binding sites were selected prioritizing on-target over off-target scores. The selected binding sites were the following: H2-018, TTTAACAAAATACTGGGAGG; H2-201, TGTATAAACATAAGGACTCT; H2-412, AGTCTTAACCCAAGGACTCC; H2-610, ATTCCTAGAGATTGTTCAAT; and H2-821, TTTACAATAAAGAGATGAAG.

To assemble RNPs, catalytically inactive *S*.*pyogenes* Cas9 (dCas9) was first expressed and purified (protocol provided upon request, but there are also alternative commercial options such as for example from New England Biolabs^®^). The crRNA and the tracrRNA with the Alt-R^®^ modifications were purchased from IDT. To ensemble RNP complexes, crRNA and the tracrRNA for each site were first pre-mixed (200 nM final in each) in the IDT Duplex Buffer, heated to 95°C for 5 min, and allowed to cool down to room temperature; then, crRNA/tracrRNA were mixed with dCas9 at 1:1 molar ratio (∼200 nM) and incubated at room temperature for 15 min. RNPs should be used freshly assembled for best results. RNPs for all binding sites were combined and mixed with 1 μg of HMW genomic DNA (1 nM final in each RNP) in 50 μL CutSmart Buffer from New England BioLabs (Cat. #B7204) and incubated at 37°C for 15 min. Genomic DNA complexed with RNPs was then digested with 40 U of Exonuclease I (Cat. #X8010; Enzymatics^®^), 100 U of Exonuclease III (Cat. #M0206L; New England Biolabs), and 20 U of Lambda exonuclease (Cat. #M0262S; New England Biolabs) simultaneously at 37°C for 9 hours in CutSmart Buffer. Exonucleases were inactivated at 80°C for 20 min. Finally, digested DNA was purified by the magnetic SPRI method (0.5x beads volume) and eluted in 25 μL 0.1x TE buffer. We used 15 μL for TELL-Seq library preparation.

### Target amplification using long-range PCR

We followed PCR conditions previously established to amplify the 13kb *SCN10A* amplicon and used the same conditions to also amplify three non-overlapping internal shorter regions: 3.9 kb, 4.1 kb, and 2.7 kb ^44^. We failed to obtain PCR product from a fourth region that would have completed the full 13 kb region in shorter amplicons, but it was ultimately not necessary for our validation purposes. PCR reactions were conducted using NA12878 genomic DNA as template. For 13 kb *SCN10A* PCR we used Supreme NZYLong DNA Polymerase (Cat. #MB331, NZYTech^®^) according to the manufacturer’s recommendations. Cycling protocol: 94°C for 5 min; 30 cycles of 94°C for 20 sec, 68°C for 30 sec, and 68°C for 14 min; final step at 68°C for 21 min. Shorter *SCN10A* amplicons were generated with Phusion Polymerase (Cat. #F565L, ThermoFisher Scientific^®^) according to the manufacturer’s recommendations. Cycling protocol: 98°C for 30 sec; 30 cycles of 98°C for 8 sec and 72°C for 90 sec; final step at 72°C for 8 min. All PCR products were processed with ExoSap-IT (Cat. #78200.200.UL; ThermoFisher Scientific) to remove primers plus one or two rounds of 0.41x HighPrep™ PCR Clean-up beads (Cat. #AC-60050; MagBio Genomics^®^) for 13 kb *SCN10A* amplicons. We used modified HighPrep™ PCR Clean-up beads procedures to remove large DNA molecules detected in the gel wells with shorter *SCN10A* amplicons, as shown in **Suppl. Fig. 5B**. Specifically, the samples were diluted to 100 μL and 41 μL of beads were next added. After a 5 min incubation at room temperature, the tubes were placed on a magnetic stand. After a minute, the 141 μL of supernatant were pipetted into a new tube, and additional 60 μL of beads were added to it before proceeding with washes and elution as specified by the manufacturer. Although there was enough PCR product to proceed to library preparation, we warn that our specific clean-up procedure led to a high product loss; thus, we recommend trying alternative bead ratios for a better recovery.

Likewise, we followed PCR conditions previously established to amplify the 3.4 kb *PIK3CA* amplicon and used the same conditions to amplify the 1.8 kb *PIK3CA* amplicon ^46^. HCC202 cells were used as cDNA source for the generation of *PIK3CA* amplicons. RNA was extracted using the *Quick*-RNA Miniprep Kit™ (Cat. #R1054; Zymo Research®) following the manufacturer’s recommendations. For cDNA generation, 1.5ug of RNA was processed using the SuperScript™ III First-Strand Synthesis System (Cat. #18080051; ThermoFisher Scientific) and either 1 uL of random hexamers or 1 uL of oligodT primer, as indicated in the Results section. PCR reactions were conducted using Phusion Hot Start II High-Fidelity polymerase (Cat. #F-565L; Thermo-Fisher Scientific) and 1 uL of stock or 1:20 dilution cDNA, as indicated in **Suppl. Fig. 5B**. Cycling protocol: 98°C for 30 sec; 30 cycles of 98°C for 10 sec, 65°C for 20 sec, and 72°C for 1 min; final step at 72°C for 8 min. PCR products were cleaned up with ExoSap-IT (Cat. #78200.200.UL; Thermo-Fisher Scientific).

All PCR products were assessed by gel electrophoresis and quantified using the Qubit™ dsDNA HS Assay kit (Cat. #Q32854; Thermo-Fisher Scientific). Primers were synthesized by Integrated DNA Technologies® and diluted to 10uM in 0.1x TE buffer. Primer sequences (5’->3’): 13kb *SCN10A* (forward) GCCATGACCATTGTTATTTGTCCAGA and (reverse) CCTGAAGAAATGTCACGGCTTGTTAG ^44^; 3.9kb *SCN10A* (forward) CACTTTGCACGAAGTGCTTG and (reverse) GCCCACACACCTCTCTTCAT; 4.1kb *SCN10A* (forward) GTGTGGGCTCTTGCTCTCAT and (reverse) GAGGTGGGAGGATGACTTGA; 2.7kb *SCN10A* (forward) TGTAATTTCTGCAGCCACGA and (reverse) CACTGGTTTCCCATTGCTCT; 3.4 kb (cDNA) *PIK3CA* (forward) TGGGACCCGATGCGGTTA and (reverse) AATCGGTCTTTGCCTGCTGA ^46^; 1.8kb (cDNA) *PIK3CA* (forward) CAGACGCATTTCCACAGCTA and (reverse) TGTGACGATCTCCAATTCCCA.

### Targeted TELL-seq protocol with Cas9-based enriched (impure) targets

Linked-read libraries were generated with an adapted version of the WGS TELL-seq Library Prep kit described here (Standard Bundle, Cat# 100035, 100036, 100003, 100004; Universal Sequencing Technology Corp.) following the Library Prep User Guide (version 8.0) and the amounts and volumes validated here for targeted phasing: 100 pg of HLS-CATCH-enriched single targets, 270 pf of a five-target HLS-CATCH-enriched panel or 15 μL of CaBagE-enriched four-target panel; 6 μL of TELL microbeads; 25 μL barcoding reaction at 35°C for 15 min followed by stabilization at 35°C for 30 min and the tagging/exonuclease reaction 35°C for 10 min. Following the wash steps as described according to the manufacture’s protocols with 13 amplification cycles (for the target panel) in 25 μL reactions. The PCR products were purified twice by the magnetic SPRI method (0.78x beads volume) and eluted in 25 μL low TE. The average size of the library was determined to be 447 bps by the TapeStation D1000 ScreenTape system (Agilent^®^), and the concentration was determined to be 2-4 nM by the Qubit High-sensitivity dsDNA Assay. Libraries (2×145 bp Illumina-compatible paired-end reads) were sequenced on a MiSeq^®^ instrument (Illumina) using the MiSeq reagent Micro kit v2, 300 Cycles (Cat. #MS-103-1002; Illumina).

### Targeted TELL-seq protocol with PCR products

Linked-read libraries were generated with an adapted version of the WGS TELL-seq Library Prep kit described here (Standard Bundle, Cat# 100035, 100036, 100003, 100004, Universal Sequencing Technology Corp.) following the Library Prep User Guide (version 8.0) and the amounts and volumes validated here for amplicon phasing (below). Bacterial and viral genomic (filling) DNAs were used to minimize human DNA collisions. The *Escherichia coli* DH10B (bacterial) strain was purchased from New England Biolabs (Cat. #FEREC0113), and genomic DNA was extracted using the *Quick*-DNA Miniprep Kit (Cat. #D3024; Zymo Research) following the manufacturer’s recommendations. BstI P-digested lambda phage gDNA was purchased from TaKaRa^®^ (Cat. #3402).

Amplicon TELL-Seq libraries were generated combining 20 pg of a single PCR product, 60 pg *E*.*coli* genomic DNA or 80 pg BstP I-digested lambda phage genomic DNA, and 6 μL of TELL microbeads (3 million microbeads), as indicated. When testing a target panel, we combined equal amounts of each fragment (for a total of 80 pg) and 6 μL of TELL microbeads (without adding filling DNA. In all tests, we used approximately 75,000 TELL microbeads for indexing (taking 1/40^th^ of the processed bead solution; and libraries were amplified with 18-21 PCR cycles. Libraries were sequenced on a MiSeq instrument (Illumina) using the MiSeq reagent Micro kit v2, 300 Cycles (Cat. #MS-103-1002; Illumina), 2×145 bp Illumina-compatible paired-end reads.

### Linked-read data analysis

Sample demultiplexing, QC reporting, and read processing of all targeted TELL-Seq and WGS TELL-Seq data were performed with TELL-Read. Mapping, variant calling, and phasing were performed using TELL-Sort, which incorporates BWA-MEM for read alignment, GATK-v4.0.3.0/HaplotypeCaller for variant calling (Broad Institute), and HapCUT2 (github/Vibansal) for phasing. The human genome (hg38) was used as reference for the analysis of all human data, except for *PIK3CA* that we used the target sequence as reference. Allele frequency threshold was set to 0.1. For the characterization of DNA collisions with lambda phage DNA, fourteen non-overlapping fragments resulting from BstP I digestion were used as reference. All software is freely available at https://www.universalsequencing.com/software. Illumina data from TELL-seq libraries were pre-converted to 10X Genomics-format. TELL-seq linked-read data was also analyzed with LongRanger v2.2.2 using the GRCh38-2.1.0 draft as reference.

### Data visualization on IGV

The Integrative Genomics Viewer, IGV-v2.11.1 (Broad Institute) was used for data visualization assisted with WhatsHap-v1.1 (github/whatshap) and it is now part of the software package that can be downloaded from the UST’s website. It is based on TELL-Sort output files, which display read mapping profiles, phased alleles, and targeted haplotypes. Visualization was also done using 10X Genomics Loupe Browser, which displays read mapping profiles, phased alleles, targeted haplotypes, and linked-read density plots, similar to those presented in IGV.

### Analysis of site behaviors for Cas9-enriched targets

Several pieces of customized scripts have been developed so that Tell-Seq reads can be aligned to the targeted regions in the reference genome for site behavior extractions from read aligned bam and phased variant vcf files. Briefly, read-aligned bam with duplicates removed from the Tell-Sort analysis has been filtered using samtools “view” program so that only the reads aligned to the targeted regions have been kept. The CIGAR string from each read in this filtered bam has been parsed so that where each nucleotide in the read been mapped to the reference has been determined. With this process, the coverage with different nucleotides on each position in the targeted region could be counted. The single nucleotide variants (SNVs) that were detected through the Tell-Sort analysis, either phased or not phased, in the vcf format, have also been mapped to the sites in the targeted regions. As a standard of the comparison, this analysis has also been applied to bam file with the reads from 10X Genomics, which is downloaded from https://github.com/genome-in-a-bottle/giab_data_indexes/blob/master/AshkenazimTrio/alignment.index.AJtrio_10Xgenomics_ChromiumGenome_GRCh37_GRCh38_06202016.HG002 (GRCh38) combined with the phased vcf file from GIAB, which is downloaded from https://ftp-trace.ncbi.nlm.nih.gov/giab/ftp/release/AshkenazimTrio/HG002_NA24385_son/latest/GRCh38/SupplementaryFiles/(HG002_GRCh38_1_22_v4.2.1_benchmark_hifiasm_v11_phasetransfer.vcf.gz)

