## Supplemental Figures for "Targeted Phasing of 2-200 Kilobase DNA Fragments with a Short-Read Sequencer and a Single-Tube Linked-Read Library Method"

A

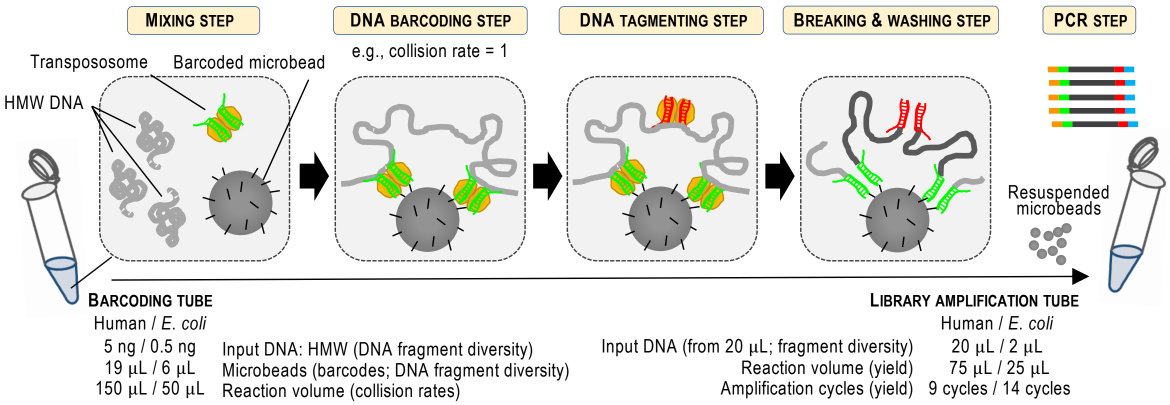


B

| TELL-Seq Applications | WGS: Animals Human Plants | Invertebrates – Metagenomics – Metagenomics – Metagenomics | Bacterial isolates | Targeted (by purity) <1-5% | Targeted (by purity) 25-100% |
|---|---|---|---|---|---|
| DNA length | 5 Gb | (decreasing) | 1 Mb | 20-300 kb | 2-30 kb |
| Input DNA amount | 5 ng | (decreasing) | 0.5 ng | 100 pg # | 20 pg #2 |
| Filling DNA* | -- | | -- | -- | 60 pg * |
| TELL-Seq reaction volume | 150 µL | (decreasing) | 50 µL | 25 µL | 25 µL |
| Reactions in standard kit | 4 | (increasing) | 12 | 12 | 12 |
| TELL microbeads | 19 µL | (decreasing) | 6 µL | 6 µL | 6 µL |
| Microbead resuspension | 20 µL | (constant) | 20 µL | 20 µL | 40 µL |
| Microbeads in PCR | 20 µL | (decreasing) | 1 µL | 2 µL | 1 µL |
| PCR volume | 75 µL | (decreasing) | 25 µL | 25 µL | 25 µL |
| PCR cycles | 9 | (increasing) | 14 | 16 | 20-21 |
| Elusion volume | 25 µL | (constant) | 25 µL | 25 µL | 15-17 µL |

### For multiple non-overlapping targets, amounts can vary between 50 pg and 100 pg per target

#2 For multiple non-overlapping amplicons, amounts vary with target number (20 ng per target for up to 4 targets; adjusted as n/80 when n>1 non-overlapping amplicons; lowest amount tested 0.04 pg)

* DNA options: BstP I-digested (14-fragment) lambda phage genome or *Escherichia coli* genomic DNA. Amounts will vary with n>1 amplicons and will be calculated after subtracting amounts of targets from the total of 80 pg. If n>4, no filling DNA will be necessary.

**Supplementary Figure 1 – TELL-Seq steps and variables**. **A.** WGS TELL-Seq workflow. Briefly, ultra-low amounts of HMW genomic DNA are mixed with transposome and TELL (barcoded) microbeads. DNA is then captured by transposome and tagged. Transposomes allow DNA recruitment on microbeads based on universal sequence homology. Microbeads provide a unique barcode to tagged DNA. A second transposome complex allows tagging an intermediate position in the barcoded fragment and breaking and washing release transposase components. Barcoded DNA is finally released and amplified to build a library for sequencing. Included key volumes, amounts, and PCR cycle numbers to process human and *E. coli* genomes, below. These conditions were used as reference to develop targeted TELL-Seq. **B.** We have adapted the WGS TELL-Seq protocol to a large diversity of genome sizes and sample complexities, which required the identification of proper amounts, volumes, and PCR cycles for each sample type (WGS Applications). Here, we propose amounts, volumes, and PCR cycles to adapt WGS TELL-Seq to samples with only one locus or a few loci (targeted TELL-Seq). In this study, we validate and further optimize these modifications with largely impure samples (<1-5% of enriched target) and relatively pure samples (25-100%). For pure samples, we hypothesized that the target should be mixed with 'competitor' non-human DNA to minimize the risk of collision, hereafter referred to as 'filling DNA', unless multiple targets were processed in the same reaction acting as filling DNA to each other (more details long this study).

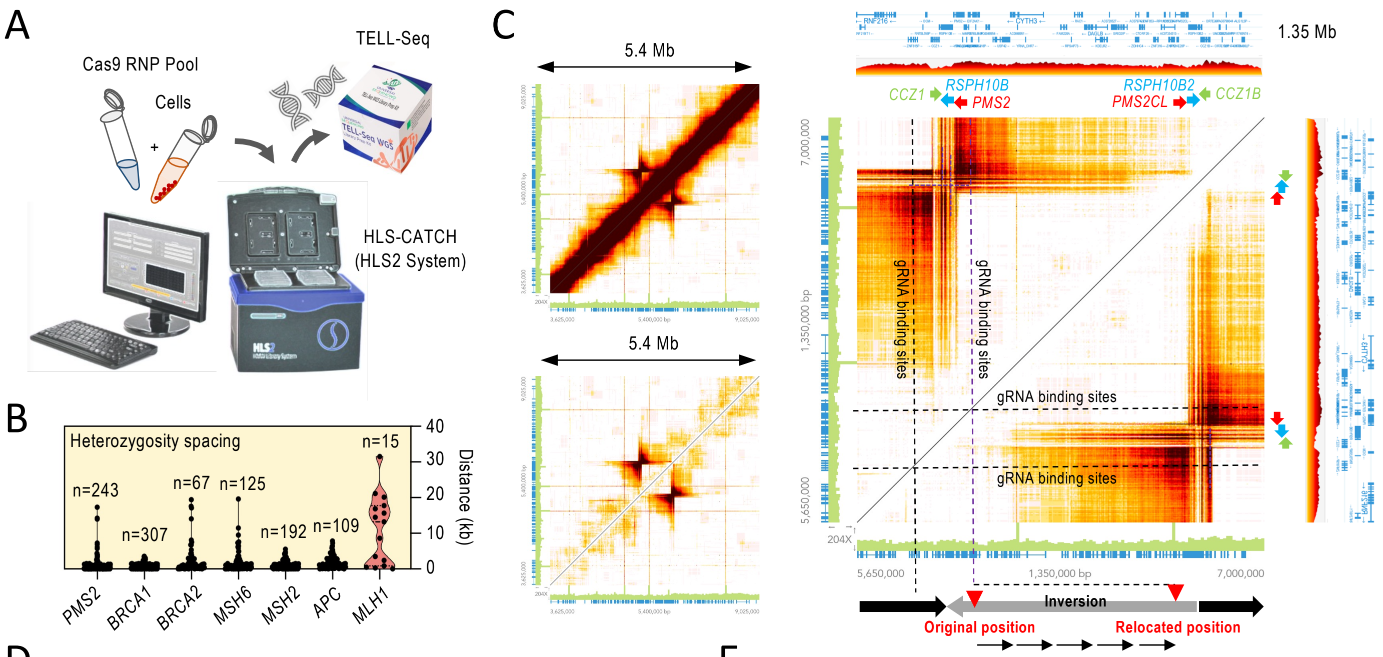


D

*MLH1*

| Chrom | Pos | NA24385_GRCh38 GIAB Reference G | A | T | C | Variant | Call | Hap1 | Hap2 | Excised MLH1 target G | A | T | C | Variant | Call | Hap1 | Hap2 |
|---|---|---|---|---|---|---|---|---|---|---|---|---|---|---|---|---|---|
| chr3 | 36923363 | 0 | 0 | 27 | 19 | C->T | het | C | T | 0 | 0 | 29 | 32 | C->T | het | C | T |
| chr3 | 36923667 | 0 | 0 | 38 | 18 | T->C | het | C | T | 1 | 0 | 28 | 25 | T->C | het | C | T |
| chr3 | 36926098 | 0 | 0 | 18 | 63 | C->T | het | T | C | 0 | 0 | 46 | 31 | C->T | het | T | C |
| chr3 | 36957616 | 0 | 30 | 0 | 32 | C->A | het | C | A | 0 | 36 | 1 | 32 | C->A | het | -- | -- |
| chr3 | 36978777 | 0 | 0 | 24 | 31 | T->C | het | C | T | 0 | 0 | 44 | 67 | T->C | het | T | C |
| chr3 | 36978786 | 30 | 0 | 0 | 23 | C->G | het | G | C | 49 | 0 | 0 | 34 | C->G | het | C | G |
| chr3 | 36991971 | 32 | 0 | 23 | 0 | T->G | het | T | G | 41 | 1 | 39 | 1 | T->G | het | -- | -- |
| chr3 | 37012077 | 48 | 34 | 0 | 0 | A->G | het | A | G | 42 | 31 | 0 | 0 | A->G | het | -- | -- |
| chr3 | 37026588 | 0 | 0 | 45 | 36 | T->C | het | T | C | 0 | 0 | 28 | 32 | T->C | het | -- | -- |
| chr3 | 37044334 | 53 | 53 | 0 | 0 | G->A | het | G | A | 15 | 27 | 0 | 0 | G->A | het | -- | -- |
| chr3 | 37061205 | 40 | 36 | 0 | 0 | G->A | het | G | A | 14 | 11 | 0 | 0 | G->A | het | -- | -- |
| chr3 | 37076854 | 43 | 36 | 0 | 0 | G->A | het | G | A | 43 | 40 | 0 | 0 | G->A | het | G | A |
| chr3 | 37080353 | 24 | 26 | 0 | 0 | G->A | het | G | A | 19 | 27 | 0 | 0 | G->A | het | G | A |
| chr3 | 37081225 | 51 | 3 | 0 | 25 | G->C | het | C | G | 39 | 0 | 0 | 35 | G->C | het | C | G |
| chr3 | 37081711 | 35 | 25 | 0 | 0 | G->A | het | A | G | 46 | 45 | 0 | 0 | G->A | het | A | G |
| chr3 | 37090324 | 25 | 0 | 41 | 0 | G->T | het | G | T | 50 | 1 | 26 | 0 | G->T | het | -- | -- |

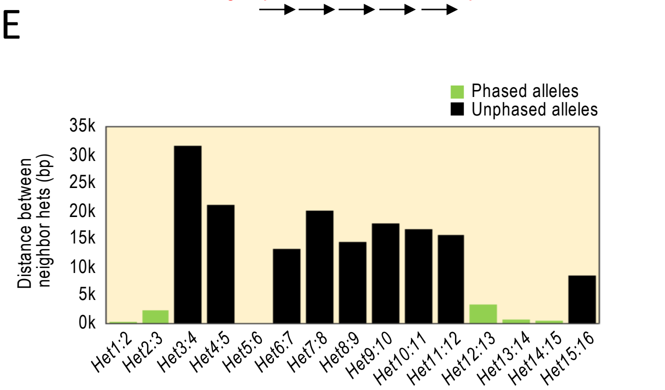


F

*PSM2*

| Chrom | Pos | NA24385_GRCh38 GIAB Reference G | A | T | C | Variant | Call | Hap1 | Hap2 | Excised PSM2 target G | A | T | C | Variant | Call | Hap1 | Hap2 |
|---|---|---|---|---|---|---|---|---|---|---|---|---|---|---|---|---|---|
| chr7 | 5990117 | 0 | 0 | 33 | 39 | C->T | het | C | T | 0 | 0 | 1 | 46 | -- | -- | -- | -- |
| chr7 | 5990656 | 50 | 25 | 0 | 0 | G->A | het | G | A | 61 | 2 | 0 | 0 | -- | -- | -- | -- |
| chr7 | 5995039 | 1 | 35 | 56 | 0 | T->A | het | T | A | 0 | 6 | 48 | 0 | T->A | het | -- | -- |
| chr7 | 5995187 | 32 | 52 | 1 | 0 | A->G | het | A | G | 2 | 51 | 0 | 0 | -- | -- | -- | -- |
| chr7 | 5995188 | 0 | 52 | 0 | 32 | A->C | het | A | C | 0 | 51 | 0 | 2 | -- | -- | -- | -- |
| chr7 | 5995797 | 0 | 39 | 0 | 36 | A->C | het | A | C | 0 | 70 | 0 | 0 | -- | -- | -- | -- |
| chr7 | 5996313 | 37 | 29 | 0 | 1 | A->G | het | A | G | 4 | 47 | 0 | 1 | -- | -- | -- | -- |
| chr7 | 5997349 | 33 | 0 | 0 | 23 | G->C | het | G | C | 58 | 0 | 0 | 1 | -- | -- | -- | -- |
| chr7 | 5998414 | 2 | 0 | 36 | 34 | T->C | het | T | C | 0 | 0 | 13 | 0 | -- | -- | -- | -- |
| chr7 | 5999091 | 0 | 1 | 23 | 25 | T->C | het | T | C | 0 | 1 | 29 | 2 | -- | -- | -- | -- |
| chr7 | 5999498 | 0 | 0 | 24 | 42 | T->C | het | T | C | 0 | 0 | 62 | 1 | -- | -- | -- | -- |
| chr7 | 6001000 | 0 | 43 | 0 | 18 | A->C | het | A | C | 0 | 57 | 0 | 1 | -- | -- | -- | -- |
| chr7 | 6002187 | 40 | 20 | 0 | 1 | G->A | het | G | A | 50 | 3 | 1 | 0 | -- | -- | -- | -- |
| chr7 | 6002205 | 0 | 18 | 44 | 0 | T->A | het | T | A | 0 | 3 | 52 | 0 | -- | -- | -- | -- |
| chr7 | 6002891 | 0 | 0 | 25 | 35 | C->T | het | C | T | 0 | 0 | 1 | 61 | -- | -- | -- | -- |
| chr7 | 6003481 | 1 | 1 | 36 | 31 | C->T | het | C | T | 0 | 0 | 2 | 45 | -- | -- | -- | -- |
| chr7 | 6004884 | 0 | 0 | 28 | 24 | C->T | het | C | T | 0 | 0 | 0 | 25 | -- | -- | -- | -- |
| chr7 | 6005019 | 0 | 0 | 37 | 30 | T->C | het | T | C | 0 | 0 | 45 | 6 | T->C | het | -- | -- |
| chr7 | 6005754 | 0 | 0 | 35 | 39 | T->C | het | T | C | 0 | 0 | 58 | 3 | -- | -- | -- | -- |

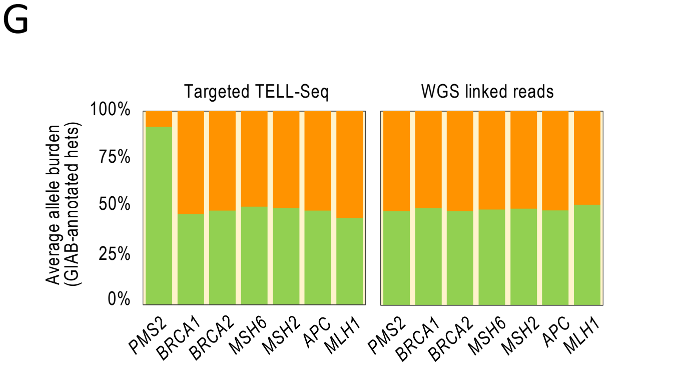


**Supplementary Figure 2 – Singularities in the *MLH1* and *PMS2* loci in HG002. A.** HLS-CATCH system (Sage Science). **B.** Violin plots showing spacing between adjacent GIAB-annotated heterozygous sites across the seven targets: *MLH1*, n=15 (datapoints), *APC*, n= 109; *MSH2*, n=192; *MSH6*, n=125; *BRCA2*, n=67; *BRCA1*, n=307; *PMS2*, n=243. **C.** WGS 10x Genomics linked-read data visualized with LongRanger Loupe (10x Genomics) focusing on the *PMS2* locus. Biding sites for gRNAs in the HLS-CATCH system are indicated. Data source: Human Pangenome Reference Consortium (https://github.com/human-pangenomics/HG002_Data_Freeze_v1.0). (*Small heatmaps*) GRCh38 chr7:3,625,000-9,025,000. Local signal (diagonal) masked in the bottom heatmap for better visualization. (*Large heatmap*) Genomic coordinates, GRCh38 chr7:5,650,000-7,000,000. Scheme of the inversion shown at the bottom. Signs of the inversion that relocates the 3' gRNA sites ~700 bp away (red arrowheads) from its position in the reference genome (purple dashed vertical line). The inversion, recurrent in the human population according to Porubsky et al., 2022, is flanked by long segmental duplications consisting in the *CCZ1*, *RSPH10B*, and *PMS2* genes and the *PMS2CL* pseudogene and the *RSPH10B2* and *CCZ1B* genes with inverted orientation in the reference genome. **D and G.** Read behaviors across a representative regions in the *MLH1* (D) and *PMS2* (G) targets to show correct genotyping but incomplete phasing (squared cells) and heterozygosity loss (incomplete genotyping and phasing) affecting Hap2 (squared cells), respectively. Left columns represent GIAB data (NA24385), right columns represent TELL-Seq data. Counts per positions are indicated by nucleotide G, A, T, C columns). Variants calls are also indicated (Variant columns). **E.** Distances between the annotated neighbor heterozygous sites. Green columns represent phased sites. Black columns represent unphased sites. **F.** Average read counts (allele burden) by target relative to the total read count. Sites selected based on GIAB annotated haplotypes.

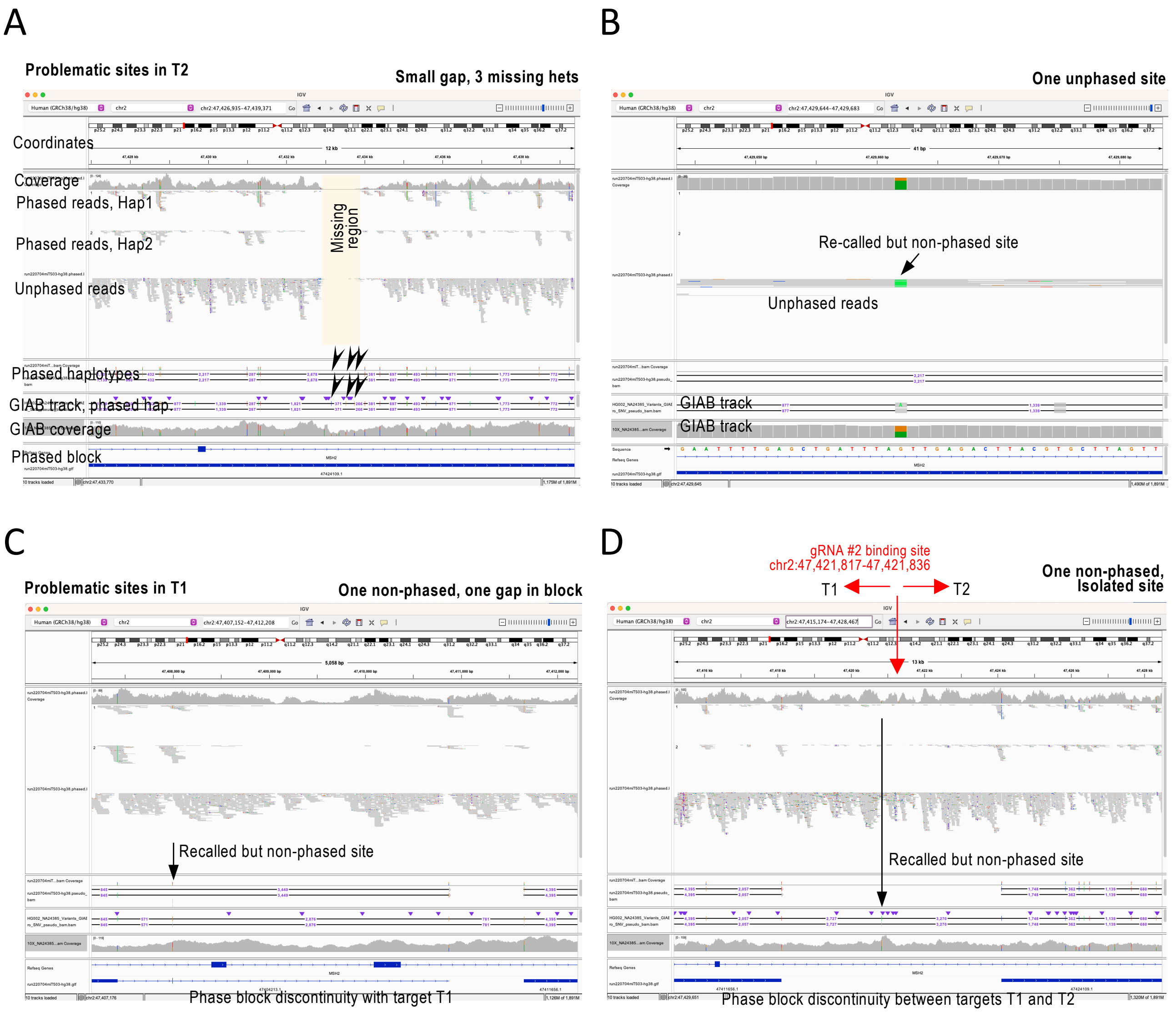


**Supplementary Figure 3 – Problematic sites in T2 and T1 targets**. Screenshots from IGV portal showing TELL-Seq data from the four *MSH2* target experiment (T1-T4). Tracks are labeled on the first screenshot for all screenshots. Phase block shown at the bottom as a blue bar. **A and B.** In T2, three heterozygous sites (indicated with arrows) were missed, explained by the presence of a small gap in the HG002 cells used in this study (in A), and one heterozygous sites was correctly re-called by not phased (shows as part of the bulk of unphased reads) (in B). **C and D.** In T1, one site was correctly re-called but not phased and there was a discontinuity in the phase block likely as the two underlying sites were not phased together (in C). Another correctly re-called heterozygous site was not phased, located at the end of the target and relatively isolated (in D).

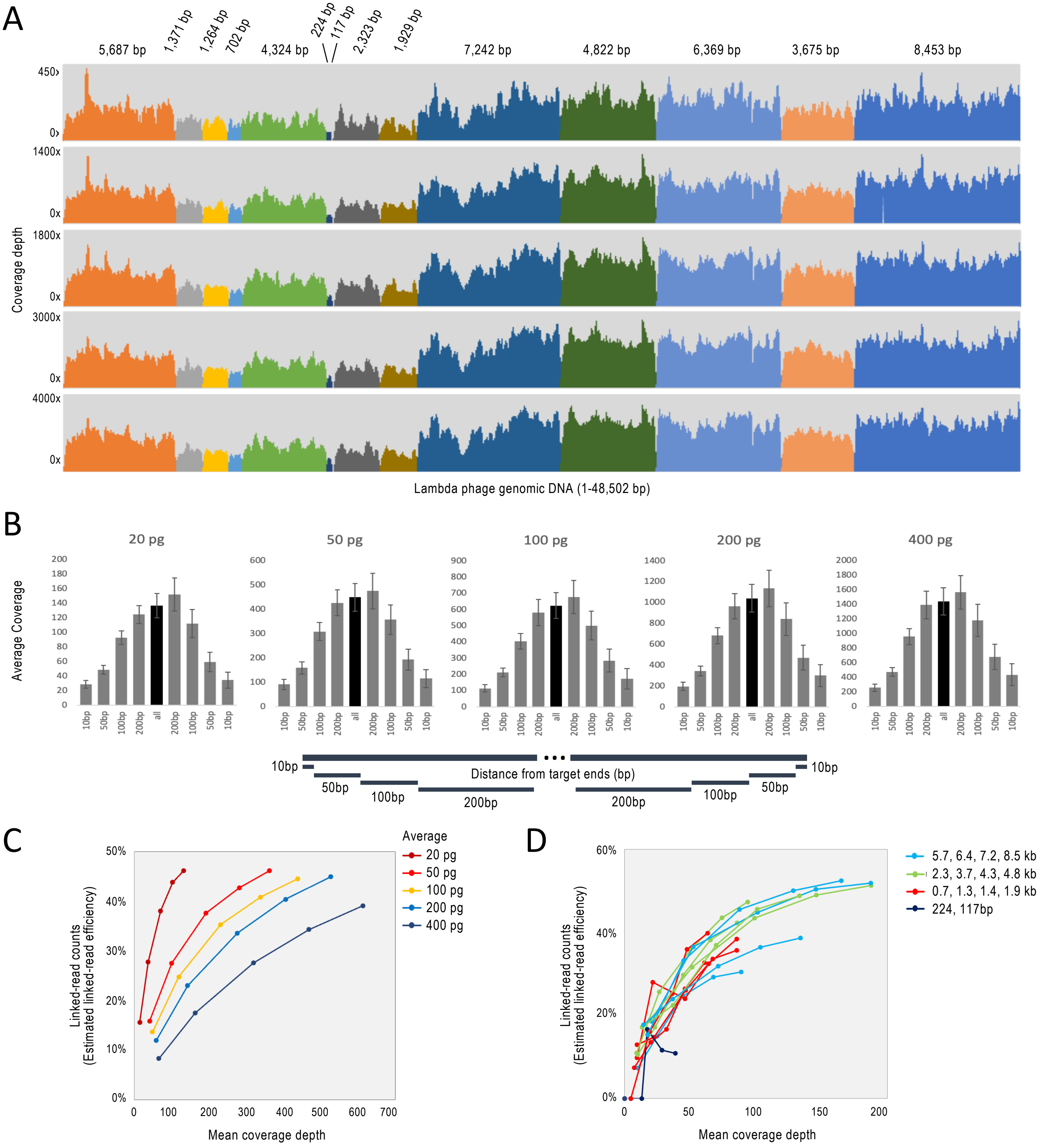


**Supplementary Figure 4 – TELL-Seq analysis of BstP I-digested lambda phage genome**. **A.** Coverage across fragments at nucleotide resolution. Fragment sizes indicated on top. **B.** Average coverage in selected regions by input amounts: 5' terminal 10 bp, 5' 11bp to 50 bp, 5' terminal 51 bp to 100 bp, 5' terminal 101 to 200 bp, rest of the fragment, 3' terminal 101 to 200 bp, 3' terminal 100 bp to 51 bp, 3' 50 bp to 11 bp, 3' terminal 10 bp. Scheme shown at the bottom. Data represents average of all 13 fragments (excluding 117 bp). Error bars represent S.E.M. **C** and **D**. Linked-read efficiencies by input after subsampling for the average from all fragments (as a %) in C. Subsampling (100%, 75%, 50%, 25%, 10%). And linked-read efficiencies by fragment size in the 20 pg sample.

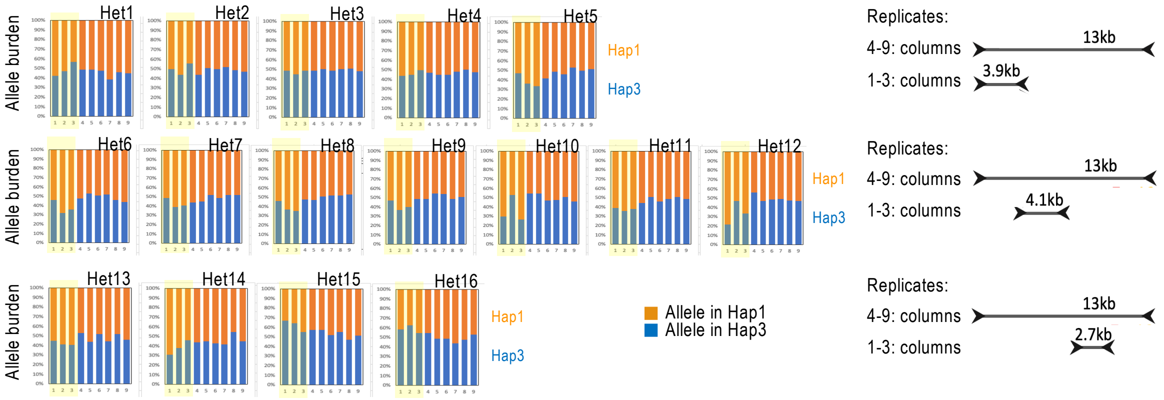


**Supplemental Figure 5 – Amplicon phasing**. Allele burden for 16 heterozygous sites in HG001 comparing multiple PCR products (three replicates for short *SCN10A* amplicons and six replicates for 13 kb *SCN10A* amplicons, as indicated on the right). In general, allele burdens fluctuate around 50%, indicating a balanced amplification in which maternal and paternal haplotypes are similarly amplified. Columns 1-3: short amplicon, three library replicates; columns 4-6: 13 kb fragment, three library replicates using lambda phage genome as filling DNA; columns 7-9: 13 kb fragment, three library replicates using *E. coli* genome as filling DNA.

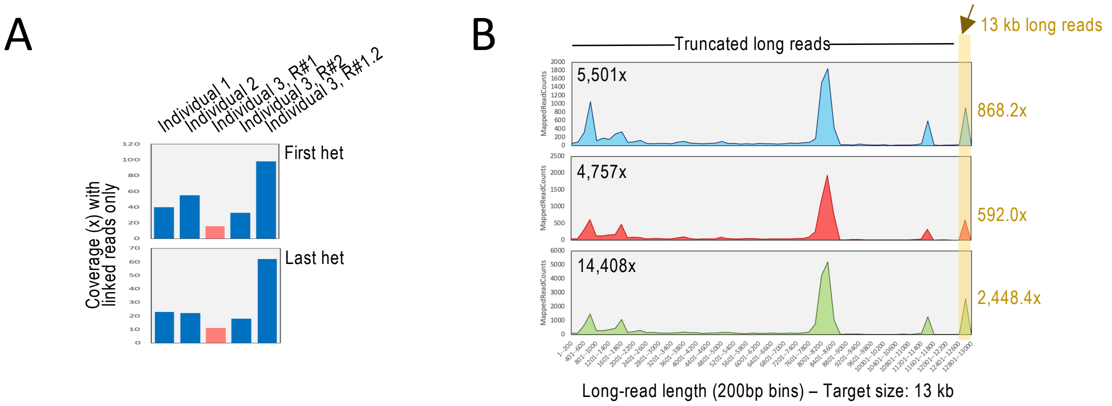


**Supplemental Figure 6 – Targeted TELL-Seq of peripheral blood-extracted genomic DNA from Hap1/Hap3 carrier individuals. A**. Linked-read coverage in the first and last heterozygous sites in the libraries shown in A. The sites with phasing issues correspond to the sites with the lowest linked-read coverage. **B**. Long-read counts (200 bp bins) in ONT data generated by Pinsach et al., 2021. Highlighted coverage from 13 kb long reads.

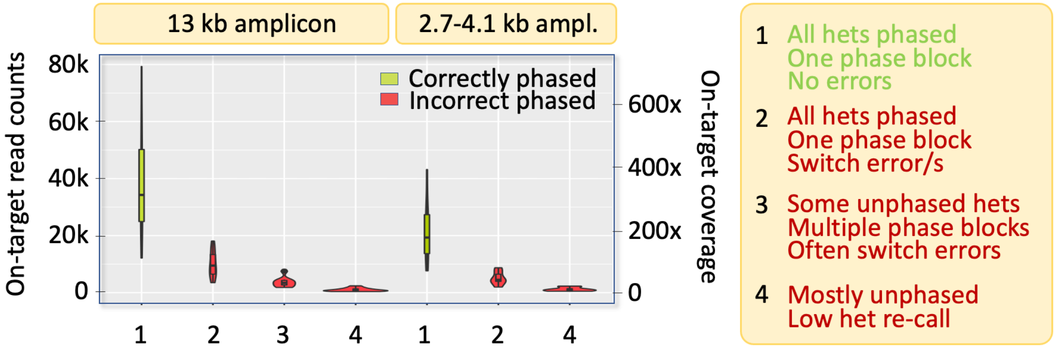


**Supplemental Figure 7 – Minimum sequencing depth for correct phasing**. Impact of sequencing depth on phasing accuracy. Violin plots containing box plots show on-target read counts and coverage (left and right axis, respectively) in three groups of data separated by phasing performance (correct phasing—i.e., all heterozygous sites are correctly recalled without phasing errors; in green; switch errors—i.e., all heterozygous sites are correctly recalled but some are incorrectly phased, in red; and missed het calls—i.e., heterozygous sites are not detected and phasing is not possible, in red. Data obtained after read duplication removal using *SCN10A* amplicons (n= 6 libraries with 13 kb and n = 3 libraries with smaller fragments) and subsampled at 50%, 25%, 12.5%, 6.25%, 3.125%, 1.56%, and 0.78% (total, n= 72 phasing).
